# Strong Positive Selection Biases Identity-By-Descent-Based Inferences of Recent Demography and Population Structure in *Plasmodium falciparum*

**DOI:** 10.1101/2023.07.14.549114

**Authors:** Bing Guo, Victor Borda, Roland Laboulaye, Michele D. Spring, Mariusz Wojnarski, Brian A. Vesely, Joana C. Silva, Norman C. Waters, Timothy D. O’Connor, Shannon Takala-Harrison

**Author notes:** These authors contributed equally to this work as joint last authors (Timothy D. O’Connor, Shannon Takala-Harrison).

## Abstract

Malaria genomic surveillance often estimates parasite genetic relatedness using metrics such as Identity-By-Decent (IBD). Yet, strong positive selection stemming from antimalarial drug resistance or other interventions may bias IBD-based estimates. In this study, we utilized simulations, a true IBD inference algorithm, and empirical datasets from different malaria transmission settings to investigate the extent of such bias and explore potential correction strategies. We analyzed whole genome sequence data generated from 640 new and 4,026 publicly available *Plasmodium falciparum* clinical isolates. Our findings demonstrated that positive selection distorts IBD distributions, leading to underestimated effective population size and blurred population structure. Additionally, we discovered that the removal of IBD peak regions partially restored the accuracy of IBD-based inferences, with this effect contingent on the population’s background genetic relatedness. Consequently, we advocate for selection correction for parasite populations undergoing strong, recent positive selection, particularly in high malaria transmission settings.

---

Malaria, a mosquito-borne disease caused by *Plasmodium* parasites, is a leading cause of illness and death in many developing countries, with an estimated 247 million cases and 619,000 malaria-related deaths in 2021 ^1^. *Plasmodium falciparum* (*Pf*) is responsible for most malaria cases and deaths. Antimalarial drugs have imposed one of the strongest selective pressures on the *Pf* genome, with the parasite having evolved resistance to nearly all drugs used as first-line therapies ^2–5^. Eastern Southeast Asia (SEA) has historically been an epicenter of emerging antimalarial drug resistance, leading to intensive malaria control efforts in this geographic region that have resulted in an 80% decrease in malaria incidence over the last twenty years ^1^. The decline in *Pf* incidence in SEA is accompanied by a structured *Pf* population, as well as decreased genetic diversity or effective population size (*N*_*e*_) ^6–9^. Monitoring the dynamics of *Pf* demography, population structure, and gene flow is a critical component of malaria surveillance efforts to inform targeted elimination intervention strategies and prevent the spread of drug-resistant parasites ^10^.

In population genetics, Identity-By-Descent (IBD) is a highly informative metric used to estimate *N*_*e*_ ^11–13^ and fine-scale population structure for recent generations ^14–17^. An IBD segment is a continuous genomic region over which a pair of isolates or genomes share an identical sequence inherited from their most recent common ancestor (MRCA) without being broken down by recombination ^18–20^. In selectively neutral scenarios, the length and positional distribution of IBD segments, as well as pairwise and population-level aggregates, can provide valuable insights into the recent evolutionary history of a population, enabling the estimation of *N*_*e*_ ^12,21^, population structure, genetic relatedness ^9^, and migration ^6,22^ in a time-specific manner ^23,24^. Although many IBD-based inference tools were initially developed for human studies, they have recently been applied to malaria parasites ^6,9,21,25,26^, despite the large differences in evolutionary parameter values between humans and *Pf* parasites. These differences include strong selection coefficients ^27^, high recombination rates (but comparable mutation rate) ^28^, and declining *N*_*e*_ ^1^ in *Pf* compared to humans. Such differences can potentially affect the quality of IBD segment detection and the patterns of IBD sharing, and thus must be considered when applying IBD-based analysis in *Pf*.

Indeed, *Pf* parasites have been under strong selection pressure due to intense malaria control efforts in recent decades, especially the widespread use of antimalarial drugs ^29–32^, resulting in the rapid emergence and spread of multidrug-resistant parasites in SEA and other malaria-endemic areas ^3^. For instance, parasites in this region have developed resistance to chloroquine, the antifolates, mefloquine, and more recently, the artemisinin derivatives and their partner drugs ^33,34^. Haplotypes harboring mutations conferring drug resistance have undergone strong selective sweeps with high selection coefficients (0.03-0.32) ^27,35^, multiple orders of magnitude greater than those usually observed in the human genome, where selection coefficients are often on the order of ∼0.001^36^. Such selective sweeps can bias inference based on IBD by changing the distribution of IBD segments and their aggregates. Positive selection increases IBD sharing at the locus under selection as well as neutral loci linked to the selected locus (genetic hitchhiking) ^37^, leading to an increase in linkage disequilibrium (LD) ^38^, long haplotypes, and long IBD segments ^37,39^. The shift of the IBD distribution has therefore been used as a signal to detect selection in both humans ^37,40^ and malaria parasite populations ^9,41^. Given the known effects of positive selection on IBD sharing patterns in human genomes ^37^, it is critical to understand whether and how IBD-based analysis is biased by positive selection in *Pf*, where selection coefficients are substantially greater than in humans.

However, the evaluation of potential selection bias in *Pf* is complicated by the high genetic relatedness of parasites in SEA, stemming from the rapid decline of the parasite population. This high background (genome-wide) genetic relatedness, LD, and IBD sharing ^6,9,42,43^ can “bury” the signal of positive selection, making it difficult to locate or correct.

Also complicating evaluation of potential selection bias is the high recombination rate observed in the *Pf* genome ^41,44,45^. Although the mutation rates in *Pf* and humans are comparable ^28^, the effective recombination rate is 60–70 times higher in *Pf* ^46^. The relatively low ratio of mutation to recombination rate in *Pf* (if ignoring background selection) leads to a small number of variants per genetic unit (low marker density), which is known to impact the ability to accurately detect IBD segments and to bias the IBD distribution ^20^. Given these factors, we need a context-specific evaluation of the effect of positive selection on IBD-based inferences of demography in *Pf*.

In this study, we employed population genetic simulations and genealogy-based true IBD segments to evaluate how positive selection, with varying parameters, affects the IBD distribution and IBD-based estimates of *N*_*e*_ and population structure in *Pf*. We proposed heuristic strategies to detect and remove genomic regions with excess IBD due to recent positive selection (IBD peaks) and evaluated whether the removal of IBD peaks mitigates positive selection-induced bias in the IBD distribution, *N*_*e*_ estimation and population structure inference. We then validated the findings from simulation analyses in empirical whole genome sequencing (WGS) datasets from low and high malaria transmission settings.

## Results

### Parasite isolates and WGS data summary

To investigate the impact of positive selection on the inference of *N*_*e*_ and population structure, we mainly focused on eastern SEA, as it has been a hotspot for drug resistance emergence ^3,47^. We analyzed WGS data from 2,055 *Pf* isolates that passed quality control and data processing filters (see Online Methods), including 751 (640 new) isolates from Cambodia and Thailand that were sequenced in-house and 1,304 eastern SEA isolates from the publicly available MalariaGEN Catalogue of Genetic Variation in *P. falciparum* v6.0 (Pf6) ^48^. The included isolates are distributed across 14 years and 18 provinces in four countries (Cambodia, Thailand, Laos, and Vietnam) (Fig. 1a). Among these isolates, 79.3%, 68.0%, and 46.1% isolates had at least 5x, 10x, and 25x coverage in >80% of the *Pf* genome, respectively (Fig. 1b). The *F*_ws_ statistic was estimated for each isolate to identify monoclonal *versus* polyclonal isolates, with 80% being classified as monoclonal isolates (*F*_ws_ > 0.95) (Fig. 1c). Among the polyclonal isolates, 44.3% harbored a predominant clone (defined in Online Methods), and the predominant haploid genome (Fig. 1d, with ratio<1.0) was included in the analyzable dataset. Isolates from West Africa (WAF) were also obtained from the MalariaGEN Pf6 database for validation of results in a high transmission setting (Fig. S1). Among WAF isolates that passed quality control, 50.7% were monoclonal, consistent with the higher multiplicity of infection (MOI) expected in a high malaria transmission setting ^28^.

**Figure 1.**
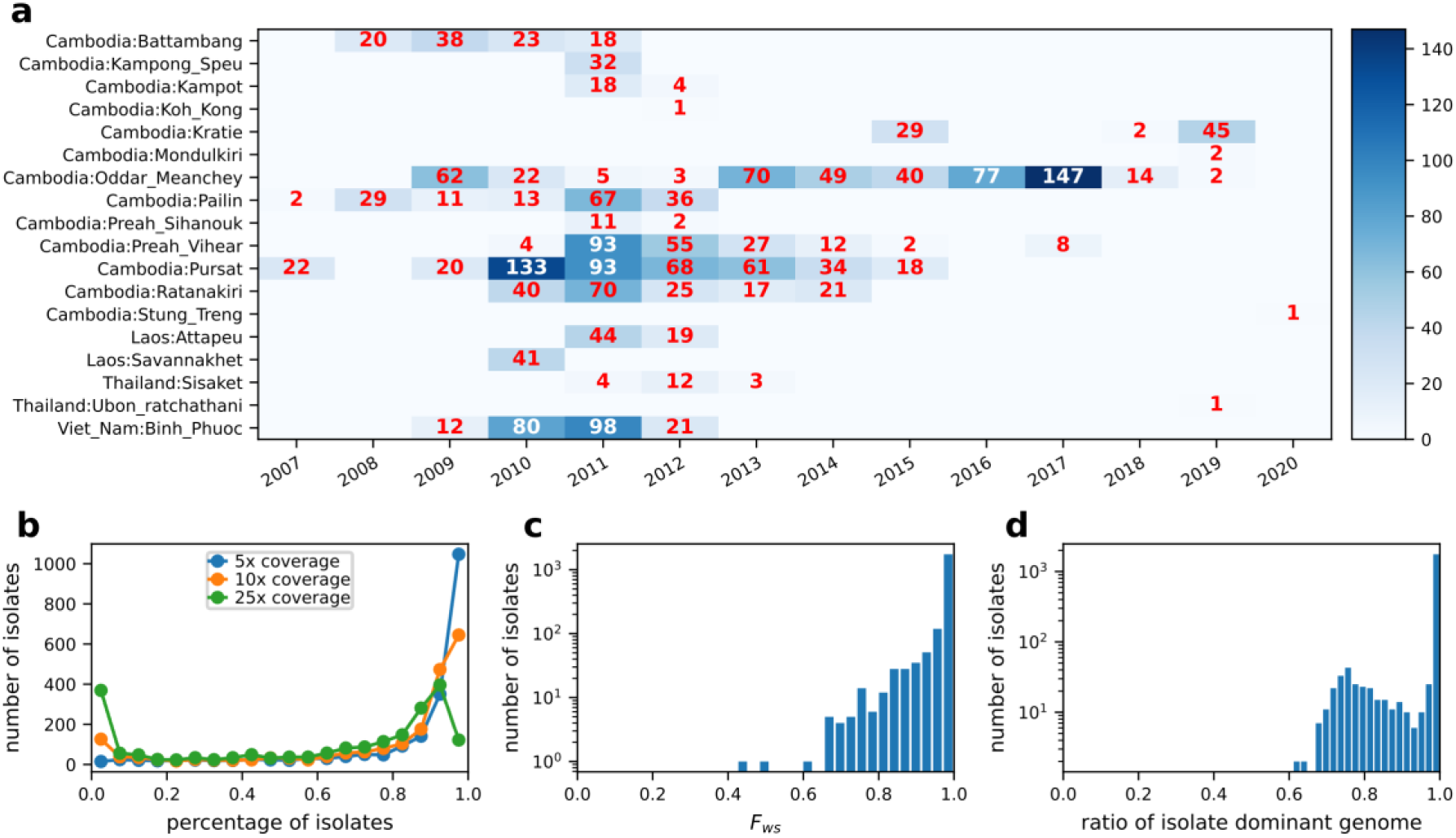
Summary of Pf parasite isolates and WGS data for Pf from SEA. **a**, Distribution of sampling location and collection year for the 2,055 analyzable samples. **b**, Distribution of genome fractions covered by at least 5, 10, and 25 sequence reads of all parasite genomes from SEA. **c**, Distribution of *F*_*ws*_ in sequenced isolates that passed quality control (genotype missingness filtering). **d**, Distribution of ratios of predominant genomes in sequenced isolates that passed quality control (determined by dEploid ^49,50^).

### Effects of positive selection on IBD distribution and IBD-based *Ne* inference

Empirical datasets sampled from a real *Pf* population often deviate from an ideal population due to various evolutionary factors, such as declining malaria incidence ^1^, parasite population structuring ^7,8,51^, selective sweeps ^3,27,34^, and asymmetrical gene flow or migration ^6^. To evaluate the direct effect of positive selection on demographic inference, we conducted population genetic simulations using simplified models that reflect parameter values observed in *Pf*, such as strong positive selection, decreasing *N*_*e*_, and a high recombination rate. We designed two categories of models: (1) a single-population model to test the effects of selection on the IBD distribution and the estimation of effective population size, and (2) a multi-population model to test the effects of positive selection on IBD-based population structure inference.

To prevent the confounding of low-quality IBD calls on the evaluation of selection effects, we implemented a true IBD inference algorithm called tskibd. The algorithm directly utilizes ancestral information from simulated (true) genealogical trees in tree sequence format ^52,53^ and avoids phased genotype-based IBD inference (Fig. S2a). We verified the quality of true IBD by comparing IBD-based *N*_*e*_ estimates (via IBDNe ^12^) with the true population size in neutral simulations under different demographic scenarios (Fig. S2b).

Different aspects of the IBD distribution represent distinct types of information about evolutionary histories, such as the time to most recent common ancestor (TMRCA, inferred based on IBD segment length) ^22,24^, genetic relatedness or population structure (inferred based on pairwise genome-wide total IBD) ^14,16,17,54^, and selection detection (inferred based on IBD positional enrichment) ^9,37,41^. We used the single population simulation model to generate genetic data under neutral (Fig. S3a) and other selection scenarios for *Pf* (Fig. S3b-d) consistent with realistic evolutionary parameters for *Pf* (see Online Methods). We found that strong positive selection impacts multiple aspects of the IBD distribution, including increasing the proportion of longer IBD segments (Fig. 2a) and isolate-pairs sharing larger genome-wide total IBD (Fig. 2b) and enriching IBD around selected sites (Fig. 2c). More importantly, we found that *N*_*e*_ (via IBDNe ^12^) is underestimated in recent generations in cases with selection compared to neutral cases (Fig. 2d), likely due to the increase in longer IBD segments (thus smaller *N*_*e*_ in more recent times).

**Figure 2.**
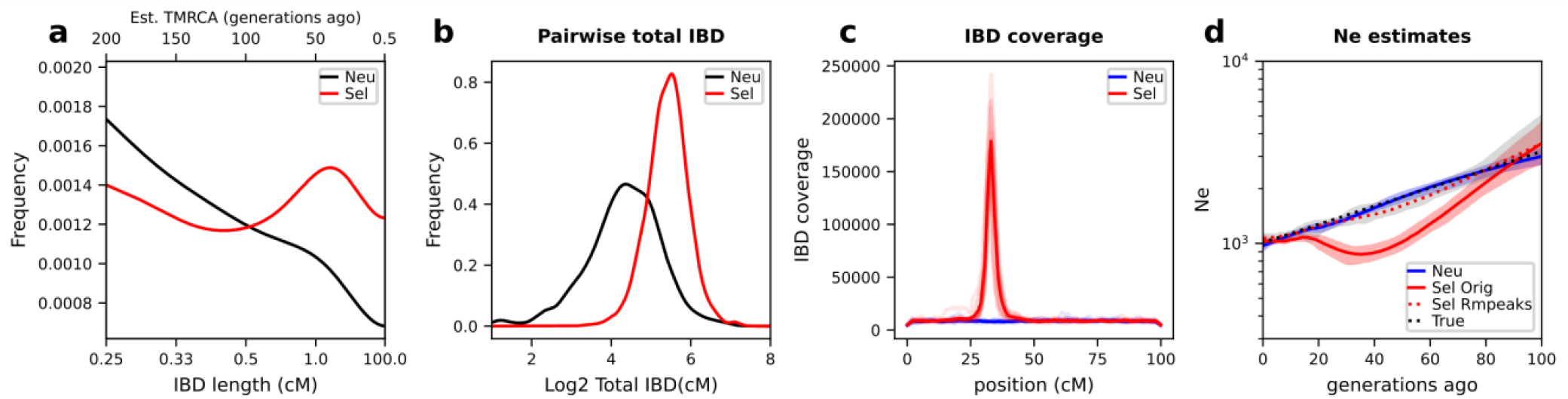
Effects of positive selection on IBD distribution and *N*_*e*_ inference. **a**-**c**, Positive selection affects various aspects of the IBD distribution, including IBD segment length (**a**), total IBD shared by a pair of isolates (**b**), and IBD location along the chromosome (**c**). Note that *x*-axis in **a** uses a custom scale for IBD length *l* (bottom) so that the estimated TMRCA (50/*l*, top) is in a linear scale. Shorter IBD segments (0.2-2 cM) were included to cover the more distant past (>25 generations ago). Lines of transparent colors in **c** represent IBD coverage for different chromosomes for the same genome set; lines of solid colors show average across chromosomes. The representative results were generated using a selection coefficient, *s*, of 0.3, a selection starting time 80 generations ago, and a single origin of the favored allele introduced at the position of 33.3 cM of each chromosome. Abbreviation: Neu, Neutral; Sel Orig, positive selection (IBD peak region not removed); Sel Rmpeaks, positive selection with IBD peak regions removed. **d**, Strong positive selection causes underestimation of *N*_*e*_ compared to neutral simulation. The difference between selection *(s* = 0.3) and neutral scenarios can be partially mitigated by removing IBD segments located within IBD peak regions. For results for different selection parameter values, see Fig. S3.

We evaluated IBD peak region identification and removal strategies to test whether positive selection-induced bias can be corrected (Fig. S4 and Online Methods). Simply put, peaks were identified on each chromosome using a threshold method, then validated through IBD-based selection statistics *X*_iR,*s*_ ^9^. Any IBD located within these validated peaks was removed. We found that removing IBD peak regions from IBD segments corrects the *N*_*e*_ estimation in the selection scenario, mimicking the neutral *N*_*e*_ estimates and true population size (Fig. 2d).

We further evaluated the impact of varying selection parameters, including selection coefficients, selection starting times (Fig. S3e-g), and the number of origins of the favored alleles (Fig. S3h-j), on IBD distribution and *N*_*e*_ estimates. In general, stronger selection (Fig. S3d), intermediate selection duration time (Fig. S3f), and a small number of origins (such as a hard sweep) (Fig. S3i) allow the establishment of the selective sweeps (the favored allele is not lost during the sweep) (Fig. S3 first column) and thus result in selection bias. Signed rank tests based on replicated simulations suggested the effects of positive selection on *N*_*e*_ estimates are statistically significant (Bonferroni-adjusted *p* values< 0.05) (Fig. S5).

### Effects of positive selection on population structure inference

Given the pronounced effects of positive selection on IBD distribution and *N*_*e*_ inference, it is vital to understand its impact on the inference of population structure. We assessed this impact using a multi-deme, one-dimensional stepping-stone model (Fig. 3a) that simulates a pattern of allele frequency gradients across subpopulations (Fig. 3b), mimicking selective sweep in a structured parasite population ^55^.

**Figure 3.**
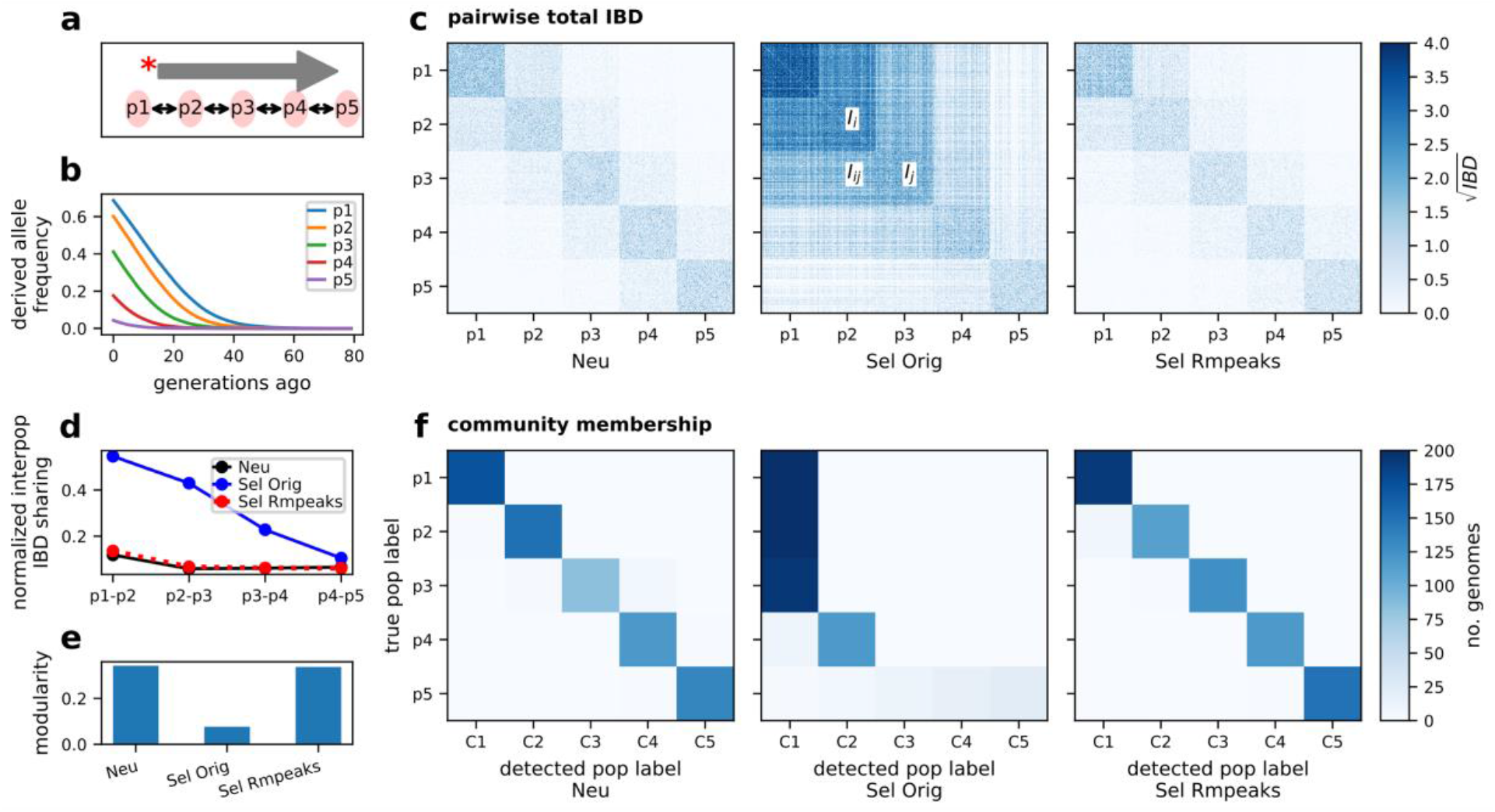
Effects of positive selection on the IBD-based population structure inference. **a**, Schematic of the one-dimensional stepping-stone model. Five subpopulations were split from an ancestral population. There is symmetrical migration between adjacent subpopulations. A favored allele was introduced into the deme from one side of the chain and spread to the other side. **b**, Average frequency trajectory of favored alleles (on each chromosome) in different subpopulations (p1 -p5). **c**, Heatmap of pairwise genome-wide total IBD under neutral, selection (s = 0.3), and selection with peaks removed. Rows and columns are ordered by true population labels. **d**, Normalized inter-population IBD sharing between nearby demes. **e**. Modularity of IBD networks with respect to the true population labels under IBD processing conditions (before and after removing IBD peaks). **f**. IBD network InfoMap community detection before (left, and middle) and after (right) removing IBD peaks. For each subplot, columns are detected community labels and rows are true population labels. The color of each block represents the number of genomes with the given true and detected communities. For results for different selection coefficients, see Fig. S6.

Under the neutral scenario with a moderate migration rate (such as 0.01, corresponding to 1% of individuals in a subpopulation being migrants from adjacent subpopulations in each generation), within-population IBD sharing dominates the pairwise sharing heatmap (Fig. 3c [left panel], and d [black line]), the total population is highly modular ^56,57^ with respect to the true subpopulation labels (Fig. 3e [left bar]), and community-detection using the InfoMap clustering algorithm ^57,58^ captures the true population structure with high consistency (Fig. 3f [left panel]).

However, with strong selection, both within- and between-population IBD sharing increases (Fig. 3c [middle panel]). This change results in an elevated ratio of inter-population to intra-population IBD sharing, (Fig. 3 d [blue line]), reduced network modularity, and collapsed community groups (Fig. 3e [middle bar] and f [middle panel], making it difficult to distinguish one population from adjacent populations. We observed similar patterns across varying selection strength (Fig. S6) and repeated simulations (Table S1), suggesting the blurring effect is selection strength-dependent.

The effect of selection on structure inference can be partially mitigated by removing IBD segments located within the genomic regions with IBD peaks (Fig. S4). After the selection correction, the dominance of within-population IBD sharing and the modularity of population is restored (Fig. 3c [right panel], d [dashed red line], and e [right bar]), and the collapsed communities become distinguishable and consistent with the true population labels (Fig. 3f [right panel]).

### Genome-wide IBD sharing and selection signals in SEA *Pf* isolates

To evaluate the effects of positive selection on IBD-based inferences in empirical *Pf* datasets, we first identified genomic regions that are under positive selection using similar methods as for simulated data, and then compared IBD-based inferences of *Pf* Ne and population structure using IBD before and after peak removal.

Previous studies have shown that IBD can be used as a metric to identify genomic regions under positive selection, including regions harboring drug-resistance mutations ^9,41^. Based on the same logic, we identified genomic regions with high IBD sharing (see Fig. S4 and Online Methods) and correlated them with known drug-resistance genes. For empirical data (without genealogical trees from simulations), we chose to use the haploid-genome-oriented HMM-based IBD caller (hmmIBD) for IBD inference ^59^. The IBD coverage profiles called by hmmIBD show peaks surrounding: (1) known drug resistance genes and genes associated with the genetic architecture of resistant parasites, such as *pfmdr1* ^60^, *pfaat1* ^41,61^, *pfcrt* ^62,63^, *dhps* ^64^, *pph* (PF3D7_1012700) ^65^, *gch1* ^66^, *kelch13* ^67–69^, and *arps10* ^65^; (2) genes related to altered sexual investment and increased transmission potential of resistant parasites, including *ap2-g*/*ap2-g2* ^43,70^ (Fig. 4a).

**Figure 4.**
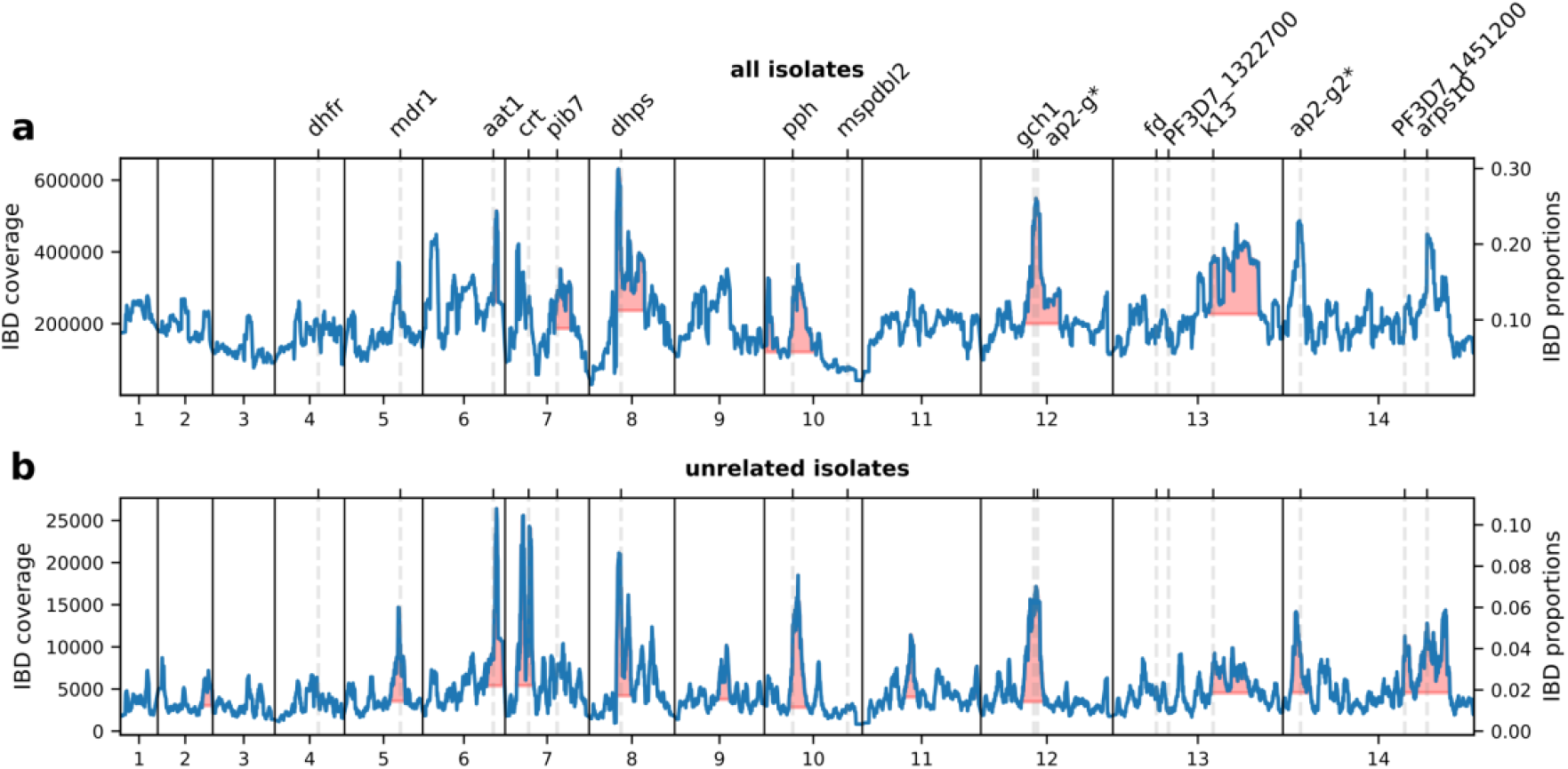
IBD coverage profile of all and unrelated *Pf* isolates in SEA. **a**. IBD coverage of all parasite genomes in SEA. Labels on the top indicate the center of known or putative drug resistance genes or genes that are under selection for sexual commitment (*). **b**. IBD coverage of unrelated genomes in SEA. Annotations in **a** are shared with **b**. Note that different scales for y axes (IBD proportions) were used to better reveal the peaks.

When including all parasite genomes, highly related and distantly related, we observe a high level of average (baseline) IBD sharing across the genome in SEA (Fig. 4a) compared to other geographic regions, such as WAF (Fig. S7), which is consistent with declines in malaria transmission owing to intensive elimination efforts in SEA ^9^. To avoid the confounding effects of high relatedness on further analysis, we applied a heuristic method to remove highly related isolates by iteratively excluding the isolate with the highest number of strong connections (defined as IBD sharing larger than half of the genome) with others. The resulting subset of isolates, hereafter called *unrelated* isolates, exhibits a five-times lower baseline IBD proportion (Fig. 4b). The low baseline IBD sharing is less noisy and more readily allows the identification of IBD peaks, including those surrounding *pfcrt* and *arps10*. (Fig. 4b). Thus, we used *unrelated* isolates and IBD peaks identified from this subset for downstream analyses.

### IBD-based inference of *N*_*e*_ and population structure in a low transmission setting with high background parasite genetic relatedness

Our simulation analyses showed that strong positive selection significantly impacts on the IBD distribution, as well as demography and population structure inference. In these simulations, removing IBD segments within the IBD peak region significantly improved the accuracy of *N*_*e*_ estimates and structure inferences. We assessed this pattern in empirical data from a low transmission setting (SEA).

First, we estimated *N*_*e*_ before removing IBD peaks. Given the high relatedness of parasite isolates in the full dataset (Fig. 4a), we focused on the unrelated isolates (*n* = 701). The *N*_*e*_ estimates based on IBD before removing the peaks via IBDNe suggest a decreasing pattern of *N*_*e*_ in SEA, from around 10^4^ to around 10^3^ in the most recent 60–80 generations (Fig. 5a, blue), consistent with a rapid decrease in malaria incidence in the last decades owing to malaria elimination efforts in this geographic region ^1^.

**Figure 5.**
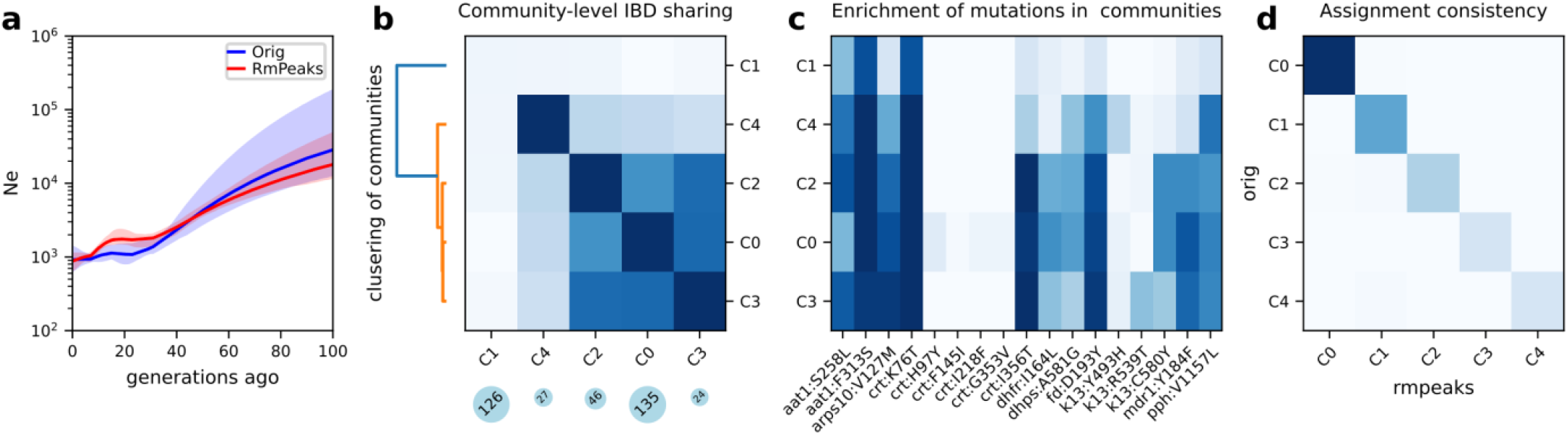
*N*_*e*_ and population structure inference in an empirical dataset from SEA. **a**, *N*_*e*_ estimates for SEA before and after removing IBD peaks. **b**, IBD network analysis of SEA data (before removing IBD peaks), including community-level IBD sharing matrix (heatmap), community size (blue circles below), and dendrogram showing hierarchical clustering of community-level IBD matrix (left). Only the largest 5 communities are plotted. **c**, Frequency of drug resistance mutations in different IBD communities. **d**, Consistency of InfoMap assignment of unrelated isolates before (x-axis) and after (y-axis) removing the peaks.

Second, we performed population structure inference before peak region removal via IBD network analyses. Using the total IBD matrix for unrelated isolates as input, we performed unsupervised community detection via InfoMap. Among the 701 unrelated isolates from SEA, we identified five communities (defined via IBD network structure analysis *instead of* geopolitical boundaries) with sizes >20 isolates. The community of the largest size, labeled as C0 (Fig. 5b) was enriched for parasites sampled from Western Cambodia (Battambang, Kratie, Oddar Meanchey, and Pursat) (Fig. S8 and Fig. S9a). In contrast, the second-largest community (C1) was comprised of isolates from a wider geographic area, including Northeastern Cambodia (Ratanakiri), Laos, and Vietnam (Fig. S8 and Fig. S9a). Isolates within C1 were distantly related given the low-average within-community IBD sharing (Fig. 5b, top left bock) compared with other communities. Hierarchical clustering of the community-level average IBD sharing matrix revealed other major communities such as C2/3/4 are closer to C0 rather than C1. The detected communities also displayed temporal dynamics. For instance, in recent years, parasites from Pursat converged into a main community including parasites from other communities; a similar pattern occurred in Oddar Meanchey 2–3–years later relative to Pursat (Fig. S9b-c). These changes are consistent with the spread of artemisinin-resistant parasite lineages from the west to the north over time.

To investigate the potential drivers of the observed population structure, we examined the distribution of non-synonymous mutations in known drug-resistance genes in the detected communities. We discovered that different communities or community groups exhibit distinct mutational landscapes at drug resistance loci (Fig. 5c). The group clustered with C0 – C0/2/3/4 – demonstrates relatively high frequencies of several resistance alleles, including those associated with artemisinin-resistance or its associated genetic background, e.g., PF3D7_0720700 C1484F, Apicoplast ribosomal protein S10 (ARPS10) V127M, PfCRT I356T, Ferredoxin D193Y, Kelch13 C493H, R539T, and C580Y, and Protein phosphatase (PPH) V1157L, as well as mutations associated with resistance to other antimalarial drugs, e.g., DHFR I164L and DHPS A581G (associated with resistance to the antifolates) and PfCRT H97Y, I218F and G353V (mutations associated with piperaquine resistance found in C0) ^8,65,71^. In contrast, the C1 communities had relatively low frequency or no mutations (including Kelch13 mutations) at these loci. The mutation landscapes for the largest 5 communities show distinct Kelch13 resistance mutation patterns, consistent with the presence of multiple artemisinin-resistant founder populations and an artemisinin susceptible population previously observed in this geographic region ^8^. These results suggest that the population structure of *Pf* in SEA is heavily influenced by drug resistance and positive selection and confirms that different founder populations harbor distinct combinations of resistant mutations ^8^.

Finally, we compared these IBD-based inferences before and after peak removal. In the SEA dataset, the removal of IBD peak regions did not significantly alter estimates of *N*_*e*_. The trajectories of *N*_*e*_ estimates are tightly overlapping with point estimates of pre-(Fig. 5a, blue) and post-segment removal (Fig. 5a, red) located within each other’s 95% confidence intervals. The inferred population structure patterns were also similar before and after the correction. The community assignments before removing IBD peaks are consistent with those when IBD peaks were not removed (Fig. 5d). Although there are some minor changes in population structure inference, the size of main communities and the community comparison metrics adjusted rand index ^72^ are largely unchanged (Table S2). The hierarchical clustering of communities shows similar grouping patterns, such as the presence of C0/2/3/4 and C1 groups (data not shown).

### Effects of removing IBD peaks on IBD-based inferences in a high transmission setting with low background parasite genetic relatedness

The effects of positive selection on IBD-based inferences observed in simulations are not corroborated by the empirical results observed in the analysis of data from parasites sampled in SEA. We hypothesized that this discrepancy stemmed from high baseline genetic relatedness observed in recent time frames in SEA (Fig. S10), despite having pruned highly-related isolates (the equivalent of first-degree relatives) (Fig. 4b). To test this hypothesis, we took two approaches: (1) we incorporated high relatedness into simulations; and (2) we evaluated an empirical dataset from a high transmission setting, WAF, where parasite relatedness is known to be lower than in SEA ^9^.

First, to incorporate high relatedness in our simulations, we condition the offspring generation process on the pedigree relatedness of parents for a subset of offspring candidates. Our results show that for *N*_*e*_ estimation, removing IBD peaks has a greater impact and provides a more accurate bias correction (resembling estimates for neutral simulation) in *unrelated* simulations (Fig. S11a) than in high-relatedness simulations (Fig. S11b). In the case of high background relatedness, peak removal results in elevated *N*_*e*_ estimates irrespective of selection and corrected estimates higher than expected under the neutral scenario, suggesting potential over-correction. For population structure inference, removing IBD peaks helps uncover hidden underlying population structures in *unrelated* simulations (Fig. S11c-d), but fails to do so in simulations under a scenario of high relatedness (Fig. S11e-f). These findings suggest that in cases with simultaneous high background relatedness and strong positive selection, the reduction in genetic diversity and the blurring in true population structure could be dominated by high background relatedness rather than by selective sweeps.

Second, to further test our hypothesis, we explored the effects of removing IBD peaks in a high malaria transmission setting by analyzing parasite isolates from WAF. Using the same criterion for unrelated isolates as in the SEA dataset, we found that removing IBD peaks in the WAF dataset, resulted in larger estimates of *N*_*e*_ for the most recent generations with non-overlapping confidence intervals around 20 generations ago (Fig. 6a). Additionally, we were able to uncover finer population structure using IBD network-based community detection algorithm after removing IBD peaks. Before selection correction, most isolates were collapsed into a dominant community (Fig. 6b). However, these isolates were assigned to many smaller communities after IBD peak removal (Fig. 6c). The change in detected communities before and after removing IBD peaks is similar to what we observe in the simulation (Fig. 3). The difference in population structure inference before and after IBD peak removal is statistically significant based on Jackknife resampling (Table S2).

**Figure 6.**
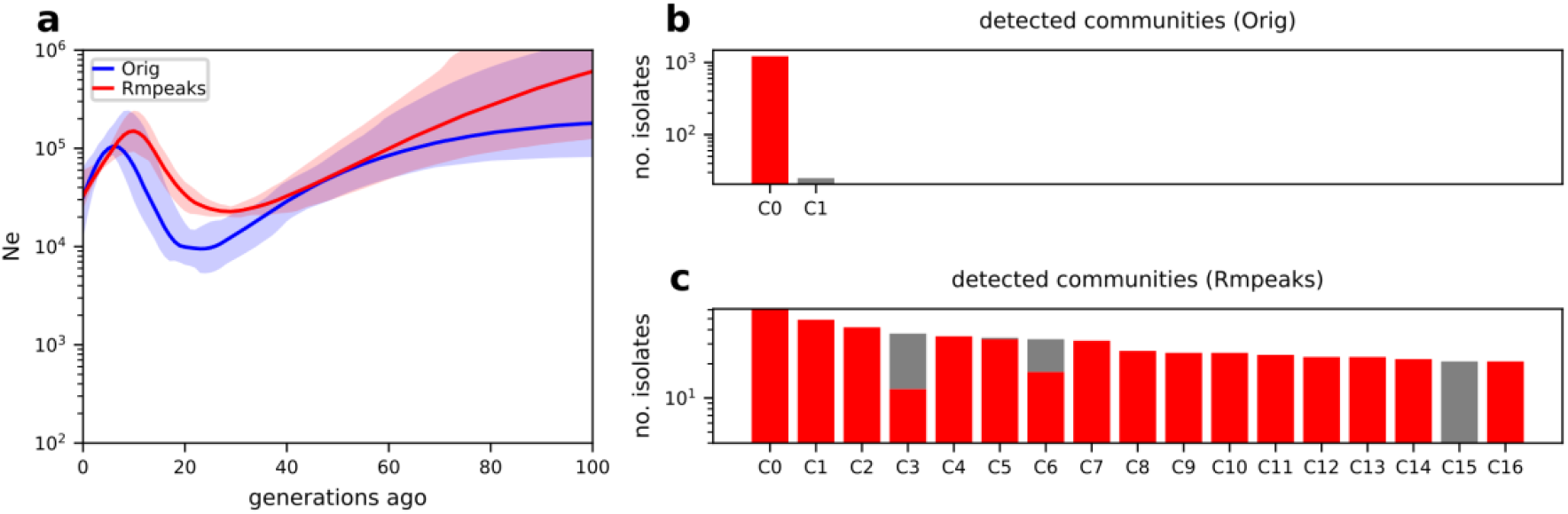
Removing IBD peaks changes the inference of *N*_*e*_ and population structure in the West African dataset. **a**, *N*_*e*_ estimates of *Pf* population in a WAF dataset before (blue) and after (red, dotted) removing peaks. **b-c**, IBD network community detection using WAF dataset before (**b**) and after (**c**) removing IBD peaks. Only communities with at least 20 isolates are shown. Removing IBD peaks allows samples of the dominant community in the original inference (**b**, the leftmost red bar) to be split into smaller communities in (**c**, red portions of many bars).

Together, these results suggest that background genetic relatedness is a key modifier of selection effects on IBD-based inference and bias correction.

## Discussion

*Pf* parasites have undergone strong and multiple selective sweeps owing to selection for resistance to antimalarial drugs, which is known to alter IBD patterns. Our simulations and analysis of field isolates demonstrated that strong positive selection can alter IBD distributions, resulting in an underestimation of *N*_*e*_ and a blurring of population structure. Our new IBD peak removal strategy can partially mitigate this bias, particularly in areas with low background relatedness (high transmission setting). Thus, we suggest excluding genomic regions under positive selection in IBD-based approaches when estimating parasite population demography in areas with high malaria transmission rates and low parasite genetic relatedness, such as sub-Saharan Africa. However, in scenarios with low transmission and high relatedness, like in SEA, correction for selection may not be necessary as such correction did not significantly change the IBD-based inferences.

Although positive selection is known to increase the likelihood of allele IBD ^9,37,40^, and accumulate longer haplotype blocks surrounding the selected sites ^39,73^, the effect of recent positive selection on the IBD segment distribution in the *Pf* genome is difficult to assess owing to the action of opposing and confounding factors, such as: (1) the strong selection that increases chromosomal local IBD sharing and thus selection bias; (2) the high genome-wide background IBD sharing (due to decreasing *N*_*e*_ and structuring of parasite population) that hides selection effects; and (3) the low SNP density (due to high recombination rate) that affects IBD call quality and the evaluation of selection effects. Our simulation approach circumvented IBD quality issues by using true IBD and mimicked high background IBD sharing by introducing high relatedness into the simulation. We provided evidence that positive selection can significantly alter IBD distributions despite the above complex context of *Pf*. Thus, ignoring selection bias in *Pf* could lead to inaccurate IBD-based inferences, particularly in high transmission settings.

The change in IBD segment length due to selection can have significant implications for time-specific inferences. This shift challenges the assumed correlation between the length of an IBD segment and its age (TMRCA) as IBD segments tend to be longer in selected regions than in neutral regions even from ancestors of similar ages. As a result, IBD-based *N*_*e*_ estimators (such as IBDNe) ^12^, time-specific migration surface estimators (such as MAPS), and other time-specific IBD analyses that rely on TMRCA estimates based on IBD segment length could all suffer from selection bias ^22,24,54^. Similarly, LD-based *N*_*e*_ estimations ^74^ are potentially biased by strong positive selection via increasing long-distance LD, which can violate the assumption that LD over longer genetic distance is informative for *N*_*e*_ of more recent time frames and shorter distance for *N*_*e*_ at distant time frames ^75,76^. Although our finding is consistent with the predicted influence of selection on the estimates of *N*_*e*_ ^76^, it contrasts with the recent finding that LD-based *N*_*e*_ estimates are virtually unaffected by natural selection ^77^. The different conclusions are likely due to the distinct ways of simulating selective sweeps. Novo *et al*. used smaller selection coefficients and did not strictly control selection loci and time (randomly chosen), which generally focuses on the relatively long-term effects of less-constrained, weak selective sweeps that mimic the human scenario; we instead conditioned on the establishment of positive selection at specific loci starting at designated time points such as 50 generations ago which mimics the recent, strong, and multi-locus selective sweeps observed in empirical *Pf* data.

The selection effect on the IBD segment length distribution can similarly impact estimates of genome-wide IBD, such as pairwise genome-wide total IBD sharing. Pairwise total IBD, similar to the fraction of sites sharing IBD, is a measure of genetic relatedness used for network-based analysis of population structure ^6,9,14,17,26,54,78,79^. As an aggregate of IBD shared across the genome, including that from neutral and non-neutral regions, pairwise total IBD-based relatedness estimates can be driven by at least two components, chromosomal local enrichment of IBD (due to non-neutral evolution), and genome-wide alteration in IBD sharing (due to changes in population structure and *N*_*e*_). Positive selection increases pairwise total IBD by increasing IBD sharing around selected loci (chromosomal local). In a panmictic malaria parasite population of large *N*_*e*_ where average IBD sharing is low (such as in Africa), pairwise total IBD is dominated by the excess chromosomal local IBD due to selection, which yields artificial patterns of relatedness unrelated to subpopulation structure. Thus, failure to correct selection bias can cause overestimation of genetic relatedness, and, further, the dominance of IBD around loci under homogeneous selection over background IBD sharing at neutral loci makes the fine-scale underlying population structure (defined by neutral regions) hard to discern.

Given the presence of selection bias, it is important to identify and correct the bias to improve the accuracy of IBD-based analyses. While there are existing heuristic methods to classify genomic loci as selected, linked neutral, or neutral regions ^80,81^, there are no specific methods to filter IBD segments for correction of positive selection-induced biases. One related exception is the IBDNe program, which internally excludes regions sharing extremely high IBD. Our peak removal strategy borrows from this idea but lowers the peak identification threshold for selected loci. We further expand the region to the chromosome median line as an attempt to include the neutral loci that are linked with the selected locus. Our work demonstrates that this peak removal method can successfully mitigate selection-induced bias in IBD-based inference of *N*_*e*_ and population structure.

As noted, the extent of positive selection bias on relatedness estimates depends on the genome-wide baseline of relatedness or IBD sharing. In a population with a high average IBD sharing across the genome (i.e., high background genetic relatedness), the pairwise total IBD is mostly driven by the genome-wide background relatedness rather than the chromosomal local increase in IBD sharing around loci under selection, making the population less likely to suffer from selection bias. Thus, background genetic relatedness is important in determining the impact of selection and the necessity for selection correction, with high transmission settings being more susceptible to selection-induced bias that requires correction. We argue against selection correction in high background relatedness scenarios, as the correction tends to be less efficient in improving the accuracy of IBD-based estimates and more likely to cause over-correction. Possible reasons for over-correction include difficulty in distinguishing selection signal (IBD peaks) from noise ^82^, and a secondary shift in IBD distribution caused by selection correction (cutting IBD segments at the peak region boundaries increases the frequency of short IBD segments). Further work is needed to understand the relationship between high background relatedness and the extent of selection bias to provide quantitative criteria on when correction is needed and how to avoid over-correction.

Besides selection bias, the application of IBD-based approaches in *Pf* research can potentially suffer from IBD call quality issues, for instance, due to relatively low SNP density (low ratio of mutation to recombination rate). Currently, IBD calling for *Pf* is performed predominantly via one of two HMM-based tools, isoRelate or hmmIBD, which report the *fraction of site IBD* (allele IBD) as the metric of genetic relatedness, with the individual IBD segments not fully evaluated or utilized ^9,59^. Our simulation models and true IBD algorithm designed for selection bias evaluation provide the foundation for development of an IBD benchmarking framework for high recombining species (ongoing work). This benchmarking framework will differ from the methods used by the top two IBD callers for *Pf*. While hmmIBD and isoRelate were validated via simple pedigree-based simulations and parent-offspring trio simulations, our framework utilizes genealogy simulation and recording (msprime/SLiM, tskit respectively) ^53,83,84^, and a tree sequence-based IBD inference algorithm (inspired by ^12^) to provide a complementary, but more flexible population-based framework for benchmarking IBD quality for different IBD callers, or evaluating IBD-based methods in non-Human species. The importance of this IBD-validation framework is also highlighted in an independent work focusing on benchmarking IBD callers for human genomes ^85^.

Despite our efforts to conduct comprehensive analyses, our present work is accompanied by several caveats. First, *Pf* evolutionary parameters are not all well-characterized and vary greatly across studies. Thus, we had to make assumptions about the realistic values of these parameters and ignore genome heterogeneity for simulation. These assumptions, if biased, might affect the accuracy of simulation analyses. Second, we simulated single-site selection simultaneously on more than one chromosome to mimic the multiple selective sweeps, but this approach may inflate the positive selection-induced bias of IBD-based estimates. In real-world scenarios, selective sweeps occur sequentially as drug policies change over time. Third, the empirical data has high heterogeneity in sampling location and time. The mixture of isolates from different years might complicate temporal interpretation of *N*_*e*_ estimates over past generations and the correspondence of IBD segment length with TMRCA. Furthermore, the presence of structure among isolates from different geographic regions as reported previously in SEA could violate the population homogeneity assumption of IBDNe ^12^ and bias *N*_*e*_ estimates. This could be partially addressed by running *N*_*e*_ estimation on isolates from a smaller time window and a specific geographic region if high variation due to a smaller sample size is tolerable. The small sample size issue could be resolved by incorporating data from the recently released, much larger MalariaGEN WGS database ^86^.

In conclusion, our study demonstrates the impact of strong positive selection on IBD-based estimates of *N*_*e*_ and population structure in *Pf*. We show that removing excess IBD within genomic regions corresponding to selective sweeps can partially correct the biases induced by positive selection and emphasizes the importance of considering selection effects when using IBD-based methods for either demography inference or population structure analysis, particularly in high malaria transmission areas. Our novel population simulation and true IBD inference framework, which employs flexible selection simulation and tree-sequenced-based IBD inference, provides a valuable tool for benchmarking IBD callers and evaluating IBD-based methods in species with extreme evolutionary parameters relative to humans.

## Acknowledgments

We would like to thank the participants in studies contributing clinical samples from which the parasite WGS data were generated. This publication uses data from the MalariaGEN Consortium and *Plasmodium falciparum* Community Project as described in “An open dataset of *Plasmodium falciparum* genome variation in 7,000 worldwide samples. MalariaGEN *et al*., Wellcome Open Research 2021642 DOI: 10.12688/wellcomeopenres.16168.1.” This work was supported by NIH 1R01AI145852 granted to ST-H and TDO by the U.S. National Institutes of Health.

## Data Availability

Raw reads of new whole genome sequence data (n=640 *Pf* isolate) will be submitted to the SRA repository (Accession numbers are currently pending). It will be publicly available after publication.

Other publicly available isolates can be found in MalariaGEN Catalogue of Genetic Variation in *P. falciparum* -v6.0, Pf6 (meta information: https://www.malariagen.net/data/catalogue-genetic-variation-p-falciparum-v6.0; raw reads: https://www.ebi.ac.uk/ena/browser/home).

## Code Availability

Source code for the true IBD inference tool *tskibd* will be publicly available before publication. Other data processing code are also available upon request.

## Material Availability

Original DNA samples are only available after discussion with corresponding authors and approval from the data collection team.

## Methods

### Parasite isolates and whole genome sequencing

The analyzed data were obtained from a publicly available repository (MalariaGEN Catalogue of Genetic Variation in *P. falciparum* - v6.0, Pf6) or from our in-house sequencing data [accession numbers pending]. The majority of analyses in this study focused on parasite isolates collected from eastern Southeast Asia (SEA). For in-house isolates, DNA was extracted from leukocyte-depleted blood samples using a Qiagen DNA Midi Kit (Qiagen, Hilden, Germany) and sequencing libraries generated using the KAPA Library Preparation Kit (Kapa Biosystems, Woburn, MA). Whole genome sequencing was performed on Illumina HiSeq 4000 or Illumina Novaseq 6000 platforms (Illumina, San Diego, CA) with generation of 150 bp paired-end reads.

We performed joint variant calling on the WGS data using a unified variant calling pipeline that follows GATK best practices and the MalariaGEN Pf6 data-generating protocol ^48^. Briefly, raw reads were first mapped to the human GRCh38 reference genome to remove host reads, with the remaining reads being mapped to the Pf3D7 reference genome (PlasmoDB_44). Mapped reads were processed using GATK MarkDuplicates and BaseRecalibrator tools, after which analysis-ready mapped reads for each isolate were used to generate per-sample calls (HaplotypeCaller /GVCF mode). These per-sample calls were combined and run through a joint-call step (GenotypeGVCFs) to obtain unfiltered multi-sample VCFs. We then used a machine learning-based variant filtration strategy, GATK VariantRecalibrator, to retain high-quality variants. Only biallelic SNPs were used in our analyses. Sites and samples were filtered based on genotype missingness and allele frequency, ensuring both per-sample and per-site missingness for filtered data were less than 0.3 and minor allele frequency ≥0.01.

Polyclonal parasite isolates were identified by calculating *F*_ws_, a metric analogous to Wright’s inbreeding coefficient, that is used to distinguish monoclonal from complex infections ^87^. Monoclonal isolates were defined as those with *F*_ws_ > 0.95 (inferred by the moimix package ^88^, and phased using dEploid ^49^, with missing genotypes imputed using Beagle 5.1 ^89^. Monoclonal isolates then served as the reference panel to extract and phase the predominant clones in polyclonal isolates, using dEploidIBD ^50^, of which the performance in predominant haplotype deconvolution has been verified by single-cell sequencing ^78^. To further ensure high haplotype quality, we used the inferred ratios of clones (haploid genomes) within an isolate to determine whether a polyclonal infection has a dominant clone: the ratio of the main clone should be greater than 0.7 and at least 3 times larger than the minor clone. The combined biallelic, phased, and imputed data, including haploid genomes from the monoclonal infections and dominant clones from the polyclonal infections, were used in downstream analyses.

### Single-population genetic data simulation

In general, simulated data were generated using forward (SLiM ^53^) and coalescent (msprime ^84^) simulators and encoded in tree sequence format ^83,90^. The advantages of combining these simulators include their efficiency (msprime), and flexibility in simulating different modes of selection (SLiM), demography, and structure. We used SLiM to perform forward-time simulations with selective sweeps of different parameter values. We then performed simplification and coalescent simulation steps ^83^. Briefly, as SLiM explicitly models each individual, the recorded tree sequences from SLiM simulation contain all explicitly modeled individuals (equal to population size N), which is more than the specified sample size. We simplify the tree sequence by subsetting so that only the requested number (sample size) of present-time genomes are kept. As the simplified tree sequences might not have fully coalesced (for instance, having multiple roots), we ran an additional backward-time simulation via msprime to fill in the top of genealogical trees, and ensure that trees coalesce into a single root, the grand common ancestor. These steps, including the forward simulation, simplification, and coalescent simulation, together generate the full genealogy ancestry for the specified samples. The full ancestry (without mutation information) was then used for two purposes: (1) to generate true IBD segments, which are elaborated below and only rely on tree topology, and (2) to allow the addition of simulated mutations via msprime onto geological tree branches and the generation of phased genotype data in variant calling format (VCF). The simulated genome consists of 14 chromosomes, each with a length of 100 centimorgans (cM), the total of which resembles a real *Pf* genome. We assume a constant recombination rate of 6.67 × 10^7^ per generation per bp (15kb/cM) ^41,44^. The mutation rate was assumed to be 1 × 10^8^ per generation per bp ^28,91^ based on which phased genotype data (VCF file) is generated. Of note, mutation information is not required for tree topology-based true IBD generation.

To evaluate the effect of selection on IBD distribution and IBD-based *N*_*e*_ inference, we simulated data using the single population model, where no population structure and migration are allowed. Under this model, we simulated genealogies with varying values of selection parameters, including selection strength, number of origins, and selection starting time. Due to genetic drift (especially for low initial allele frequency or weak selection coefficient), the favored allele (under selection) can be lost and in turn no effective selection would be observed in the present samples. To condition on the establishment of selection, we rerun the simulation with different seeds up to 100 times until the favored allele is not lost at the present-day time.

We ran simulations under different *N*_*e*_ scenarios, including constant *N*_*e*_ (see multi-population model below), and exponential decrease, where the ancestral population size was assumed to be 10,000. Our choice of *N*_*e*_ 10,000 is an intermediate value within the large range of *N*_*e*_ estimates for *Pf* in Southeast Asia from 10^3^ to 10^5 27,92,93^. For selection effect evaluation of IBD distribution and *N*_*e*_ estimates, we chose to use the exponential decreasing demographic model to mimic the *Pf* demography in SEA.

### IBD calling and removing highly related isolates

We implemented a true IBD inference algorithm (tskibd, available at: github.com/bguo068/tskibd) based on true genealogical trees, to circumvent the bias due to low-quality IBD calls from mutation information (phased genotype data), and directly test the effect of selection on IBD and IBD-based inference. The tskibd algorithm was implemented on top of the tskit C API. Our definition of IBD segments closely follows Browning *et. al*.’s work ^20^. The main concept in this algorithm is that for each pair of haploid genomes, if the most recent common ancestor (MRCA) stays the same across adjacent marginal trees so that span is longer than a specified threshold such as >2cM, this unbroken, long, and shared ancestral segment is defined as an IBD segment for this pair of haploid genomes. The algorithm produces IBD segment records for all pairs of haploid genomes, chromosome by chromosome. In addition to start and end coordinates, the produced IBD records contain additional useful information, such as whether the segment contains the favored allele (if recorded from SLiM simulation) and MRCA and its age, on which we can depend for accurate segment filtration that could not be done using other IBD callers. We apply this algorithm for all simulated data where true genealogy is already available.

When working with an empirical dataset, the true genealogy is not available. We chose to use hmmIBD for IBD calling from our empirical dataset as it was developed specifically for haploid parasites including *Pf* and has been commonly used in the malaria research community ^59,94,95^. As instructed in hmmIBD documentation, we modified the recombination rate in the source code to make it consistent with our simulations (15kb/cM) and specified the -n option to 100 to call IBD segments from MRCAs in the recent 100 generations. The input includes phased and imputed genotype data from all monoclonal samples and the dominant haploid genome from polyclonal samples (see criteria mentioned above). Tracts < 2 cM in length were excluded as these short tracts tend to lack statistical support to be confidently called IBD ^9,20^.

We used a heuristic method to remove isolates that are highly related while maximizing the number of remaining isolates. First, we defined a pair of isolates (represented by haploid genomes, either from monoclonal infections or dominant clone from polyclonal infections) as *highly related* if their genome-wide (chromosome PF3D7_01_v3 to PF3D7_14_v3) IBD sharing is no less than half of the genome size. We then built an adjacency matrix by including all highly related isolates. The relatedness of each pair is either high (represented by 1) or low (represented by 0). We made a network/graph based on the above high relatedness adjacency matrix. From the graph, we iteratively deleted the node with the highest number of connections (degrees) and the node’s related connections and append the deleted node to a list until no connection was present in the graph. The list of deleted nodes (from the high relatedness network) is the subset of isolates to be removed in order to keep all remaining isolates “*unrelated*” (not highly related). Finally, we removed all IBD segments involving the to-be-deleted isolates from input IBD. IBD shared among unrelated isolates was used for *N*_*e*_ and population structure inference.

### IBD coverage profiling and peak identification and removal

We calculated IBD coverage as the number of IBD segments overlapping each position of a list of evenly spaced (0.1 cM) sampling points along each chromosome. IBD peak candidates were identified as regions with IBD sharing higher than two 5%-trimmed standard deviations above the trimmed mean (core region) which is extended on both sides until the coverage reaches the median coverages of the chromosome (extension region). To differentiate noise from real selection signals, we calculated an IBD-based positive selection statistics *X*_iR,*s*_ ^9^ for each SNP and treat each SNP with a statistically significant selection signal as a *hit*. We then overlaid IBD peak candidates and *X*_iR,*s*_-based hits. An IBD peak candidate that contains ≥ 1 hit will be defined as a verified IBD peak.

The identified and verified peak regions (core and extension regions) suggested signals of selection. For each IBD segment, we removed the whole segment if it was contained within a peak region, or removed the part of the segment that was overlapping with any peak region. The remaining segment(s) longer than 2 cM were kept. The removal of IBD peaks creates empty ranges of zero IBD coverage. When preparing input for IBDNe ^12^, we followed the internal algorithm used in IBDNe and split chromosomes into contigs by treating each contiguous region with non-zero IBD coverage as an independent contig (chromosome).

### Effective population size estimation

We used IBD segments no shorter than 2 cM as input of IBDNe. To be compatible with tools designed for diploid species, such as IBDNe for *N*_*e*_ inference, we processed the haploid-level IBD to diploid-level IBD by randomly assigning two haploid genomes (isolates) to a pseudo–heterozygous diploid.

Processed IBD segments and a constant-rate (15kb/cM) recombination map were used to estimate *N*_*e*_ over the last 150 generations using IBDNe, with a focused interpretation on the latest 100 generations ^12^. The minregion parameter was changed from 50 cM to 10 cM to include shorter *Pf* contigs/chromosomes. For selection-aware *N*_*e*_ estimation, we removed and/or split IBD segments corresponding to IBD peaks as described above. Bootstrap sampling (n=80/generation by default) was used to estimate uncertainty for empirical datasets.

### Multi-population genetic data simulation

To evaluate the effects of selection on population structure, we simulated genealogies of genomes under positive selection involving multiple subpopulations using a 1-dimensional stepping-stone model. Five subpopulations, split from the same ancestral population (*N*_*e*_ =10,000) 500 generations ago, each with an effective population size of 10,000, are connected by symmetrical migration between neighboring populations (p1 ↔ p2 ↔ p3 ↔ p4 ↔ p5, with a migration rate of 10^−5^ until selection starts). A favored allele (hard selective sweep) originates from the leftmost population (p1) 80 generations before the present, expands and spreads toward the rightmost population (p5) via migration. The selection pressure (with selection coefficients varying from 0 to 0.3) on this allele is the same across all populations (i.e., uniform selection rather than heterogeneous selection). With appropriate combinations of the selection and migration parameters (0.01 is used after selection starts), this model mimics the gradient of allele frequencies that occur when a selective sweep spreads across populations ^3^. Similar to the single population simulation model and other simulation studies, we reran each simulation up to 100 times until the present-day allele frequency in the leftmost subpopulation is at least 0.2, in order to reduce the possibility of the favored allele being lost during the sweep process, and measure effects of established positive selection.

### Population structure inference

We inferred the population structure based on a pairwise genome-wide total IBD matrix. For the empirical dataset, we used only longer segments (>4 cM) to build a total IBD matrix that reflects a more recent structure. We set elements of the total IBD matrix to zeros if they are less than 5 cM to reduce noise and reduce the density of the matrix. To test if there is an isolation-by-distance pattern, we plotted a heatmap of an unclustered IBD matrix that was ordered by sampling location. For unsupervised clustering, we combined two different approaches: (1) Run InfoMap algorithm (a community-detection method implemented in python-igraph package) on an IBD network that is built upon the square-transformed IBD matrix 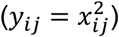. This square transformation can help better reveal the fine-scale structure within the simulated population. (2) Run an agglomerative hierarchical clustering via the mean linkage criterion (implemented in the scikit-learn python package) on a community-level average IBD matrix to assess how communities were related. The original IBD matrix was transformed via the formula **Y** = max(**X**)/(**X** − min(**X**) + *δ*), where *δ* is a small number 0.001, to convert it from a similarity matrix to a dis-similarity matrix.

Additionally, when the true population structure was known (in simulated data from the multi-population model), we calculated the (1) *modularity* of the IBD network with respect to the true structure to measure how the genomes within each true population are separated from other groups using a function in the python-igraph package, and (2) the *normalized inter-population IBD sharing* defined as the ratio of inter-population sharing *I*_*i,j*_ to the square root of the product the intra-population sharing *I*_*i*_ and *I*_*j*_. To evaluate the effect of removing IBD peaks (selection effects), the above inference or calculation was repeated on IBD data with peak regions excluded.

### Simulating high background relatedness

Incorporation of high relatedness/inbreeding into the single-population or multi-population model is implemented in the SLiM simulation script by modifying the acceptance/rejection rate of offspring candidates so that offspring from more related parents (defined by pedigree, called the coefficient of relationship ^96^) are favorably chosen as parents for the next generation. To avoid an extremely quick increase in relatedness over generations, we leave a fraction of the offspring slots without conditioning on high pedigree relatedness of their parents (thus random) and the rest conditioning on parents’ high relatedness that is not extreme, such as 1/32–1/4. For the multi-population simulation model, new migrants would be less favorably chosen as parents if only conditioning offspring candidates on the parent’s relatedness, which could lead to a reduced effective migration rate. We mitigate this by accepting all offspring candidates if one of the parents is a first-generation migrant.

### Statistical methods

For verifying IBD peaks, we calculated IBD-based statistics *X*_iR,*s*_ ^9^ for each SNP. We determined the statistical significance after the *Benjamini/Hochberg* correction using the multipletests function in the Python package statsmodels ^97^.

For IBDNe results in empirical data, the 95% confidence intervals from bootstrap sampling (within the output of IBDNe inference) were used to compare *N*_*e*_ estimates based on IBD-filtered and unfiltered inputs. For simulated data, we used the Wilcoxon signed-rank test to examine point estimates of 30 replicated simulations before and after removing IBD peaks for each generation. The *p*-values were adjusted using Bonferroni correction as estimates from nearby generations are highly correlated.

For comparing community-detection-based population assignments in empirical datasets, we calculated the *Adjusted Rand Score* ^72^ to quantify the level of consistency between memberships before and after removing IBD peaks. We used the Jackknife resampling to obtain the confidence intervals by randomly excluding one chromosome and rerunning the community detection analyses for each sampling. We defined the difference between assignments as statistically significant when the upper bound of confidence intervals (the third quartile + 1.5 interquartile range) did not contain the value 1.0, given the expected values of the Adjusted Rand Score for identical assignments are 1.0. For simulated data with known true population labels, we calculated Adjusted Rand Scores against the true population labels. To determine the uncertainty, we ran 30 replicates of simulations of the same model and parameter values but different seeds. We compared the Adjusted Rand Scores (each against the truth) before and after removing IBD peaks using the paired *t*-test.

A *p*-value, or an adjusted *p*-value if corrected, less than 0.05 is treated as statistically significant, unless otherwise indicated.

## Reference

1. World Health Organization. World malaria report 2022. (World Health Organization, 2022).

2. Ashley, E. A. et al. Spread of Artemisinin Resistance in Plasmodium falciparum Malaria. N. Engl. J. Med. 371, 411–423 (2014).

3. Hamilton, W. L. et al. Evolution and expansion of multidrug-resistant malaria in southeast Asia: A genomic epidemiology study. Lancet Infect. Dis. 19, 943–951 (2019).

4. Packard, R. M. The origins of antimalarial-drug resistance. N. Engl. J. Med. 371, 397–9 (2014).

5. Imwong, M. et al. The spread of artemisinin-resistant Plasmodium falciparum in the Greater Mekong subregion: A molecular epidemiology observational study. Lancet Infect. Dis. 17, 491–497 (2017).

6. Shetty, A. C. et al. Genomic structure and diversity of Plasmodium falciparum in Southeast Asia reveal recent parasite migration patterns. Nat. Commun. 10, 1–11 (2019).

7. Dwivedi, A. et al. Plasmodium falciparum parasite population structure and gene flow associated to anti-malarial drugs resistance in Cambodia. Malar. J. 15, 319 (2016).

8. Miotto, O. et al. Multiple populations of artemisinin-resistant Plasmodium falciparum in Cambodia. Nat. Genet. 45, 648–655 (2013).

9. Henden, L., Lee, S., Mueller, I., Barry, A. & Bahlo, M. Identity-by-descent analyses for measuring population dynamics and selection in recombining pathogens. PLoS Genet. 14, e1007279–e1007279 (2018).

10. World Health Organization. World malaria report 2018. (World Health Organization, 2018).

11. Palamara, P. F., Lencz, T., Darvasi, A. & Pe’er, I. Length Distributions of Identity by Descent Reveal Fine-Scale Demographic History. Am. J. Hum. Genet. 91, 809–822 (2012).

12. Browning, S. R. & Browning, B. L. Accurate Non-parametric Estimation of Recent Effective Population Size from Segments of Identity by Descent. Am. J. Hum. Genet. 97, 404–418 (2015).

13. Fournier, R., Reich, D. & Palamara, P. F. Haplotype-based inference of recent effective population size in modern and ancient DNA samples. Romain Fournier,David Reich, Pier Francesco Palamara. BioRxiv Prepr. Serv. Biol. (2022).

14. Han, E. et al. Clustering of 770,000 genomes reveals post-colonial population structure of North America. Nat. Commun. 8, (2017).

15. Taylor, A. R. et al. Quantifying connectivity between local Plasmodium falciparum malaria parasite populations using identity by descent. PLOS Genet. 13, e1007065 (2017).

16. Nait Saada, J. et al. Identity-by-descent detection across 487,409 British samples reveals fine scale population structure and ultrarare variant associations. Nat. Commun. 11, 6130 (2020).

17. Belbin, G. M. et al. Toward a fine-scale population health monitoring system. Cell 184, 2068-2083.e11 (2021).

18. Browning, S. R. & Browning, B. L. Identity by Descent Between Distant Relatives: Detection and Applications. Annu. Rev. Genet. 46, 617–633 (2012).

19. Ralph, P. & Coop, G. The Geography of Recent Genetic Ancestry across Europe. PLOS Biol. 11, e1001555 (2013).

20. Zhou, Y., Browning, S. R. & Browning, B. L. A Fast and Simple Method for Detecting Identity-by-Descent Segments in Large-Scale Data. Am. J. Hum. Genet. 106, 426–437 (2020).

21. Morgan, A. P. et al. Falciparum malaria from coastal Tanzania and Zanzibar remains highly connected despite effective control efforts on the archipelago. Malar. J. 19, 1–14 (2020).

22. Al-Asadi, H., Petkova, D., Stephens, M. & Novembre, J. Estimating recent migration and population-size surfaces. PLOS Genet. 15, e1007908–e1007908 (2019).

23. Browning, B. L. & Browning, S. R. A Fast, Powerful Method for Detecting Identity by Descent. Am. J. Hum. Genet. 88, 173–182 (2011).

24. Baharian, S. et al. The Great Migration and African-American Genomic Diversity. PLoS Genet. 12, 1–27 (2016).

25. Taylor, A. R., Jacob, P. E., Neafsey, D. E. & Buckee, C. O. Estimating relatedness between malaria parasites. Genetics 212, 1337–1351 (2019).

26. Taylor, A. R., Echeverry, D. F., Anderson, T. J. C., Neafsey, D. E. & Buckee, C. O. Identity-by-descent with uncertainty characterises connectivity of Plasmodium falciparum populations on the Colombian-Pacific coast. PLOS Genet. 16, e1009101 (2020).

27. Anderson, T. J. C. et al. Population Parameters Underlying an Ongoing Soft Sweep in Southeast Asian Malaria Parasites. Mol. Biol. Evol. 34, 131–144 (2017).

28. Camponovo, F., Buckee, C. O. & Taylor, A. R. Measurably recombining malaria parasites. Trends Parasitol. (2022) doi:10.1016/j.pt.2022.11.002.

29. Wootton, J. C. et al. Genetic diversity and chloroquine selective sweeps in Plasmodium falciparum. Nature 418, 320–323 (2002).

30. Bloland, P. B., Surveillance, W. H. Organization. A.-I. D. R. & Team, C. Drug resistance in malaria / peter B. Bloland. A background document for the WHO global strategy for containment of antimicrobial resistance (2001).

31. White, N. J. Antimalarial drug resistance. J. Clin. Invest. 113, 1084–1092 (2004).

32. Greenwood, B. Anti-malarial drugs and the prevention of malaria in the population of malaria endemic areas. Malar. J. 9, S2 (2010).

33. Ménard, D. & Fidock, D. A. Accelerated evolution and spread of multidrug-resistant Plasmodium falciparum takes down the latest first-line antimalarial drug in southeast Asia. Lancet Infect. Dis. 19, 916–917 (2019).

34. Blasco, B., Leroy, D. & Fidock, D. A. Antimalarial drug resistance: Linking Plasmodium falciparum parasite biology to the clinic. Nat. Med. 23, 917–928 (2017).

35. Anderson, T. J. C. Mapping drug resistance genes in Plasmodium falciparum by genome-wide association. Curr. Drug Targets Infect. Disord. 4, 65–78 (2004).

36. Peyrégne, S., Boyle, M. J., Dannemann, M. & Prüfer, K. Detecting ancient positive selection in humans using extended lineage sorting. Genome Res. 27, 1563–1572 (2017).

37. Albrechtsen, A., Moltke, I. & Nielsen, R. Natural Selection and the Distribution of Identity-by-Descent in the Human Genome. Genetics 186, 295–308 (2010).

38. Stephan, W., Song, Y. S. & Langley, C. H. The Hitchhiking Effect on Linkage Disequilibrium Between Linked Neutral Loci. Genetics 172, 2647–2663 (2006).

39. Sabeti, P. C. et al. Detecting recent positive selection in the human genome from haplotype structure. Nature 419, 832–837 (2002).

40. Browning, S. R. & Browning, B. L. Probabilistic Estimation of Identity by Descent Segment Endpoints and Detection of Recent Selection. Am. J. Hum. Genet. 107, 895–910 (2020).

41. Amambua-Ngwa, A. et al. Major subpopulations of Plasmodium falciparum in sub-Saharan Africa. Science 365, 813–816 (2019).

42. Cerqueira, G. C. et al. Longitudinal genomic surveillance of Plasmodium falciparum malaria parasites reveals complex genomic architecture of emerging artemisinin resistance. Genome Biol. 18, 78 (2017).

43. Early, A. M. et al. Declines in prevalence alter the optimal level of sexual investment for the malaria parasite Plasmodium falciparum. Proc. Natl. Acad. Sci. 119, e2122165119 (2022).

44. Conway, D. J. et al. High recombination rate in natural populations of Plasmodium falciparum. Proc. Natl. Acad. Sci. U. S. A. 96, 4506–4511 (1999).

45. Kong, A. et al. A high-resolution recombination map of the human genome. Nat. Genet. 31, 241–247 (2002).

46. Venter, J. C. et al. The Sequence of the Human Genome. Science 291, 1304–1351 (2001).

47. Wongsrichanalai, C. & Meshnick, S. R. Declining Artesunate-Mefloquine Efficacy against Falciparum Malaria on the Cambodia– Thailand Border. Emerg. Infect. Dis. 14, 716–719 (2008).

48. MalariaGEN et al. An open dataset of Plasmodium falciparum genome variation in 7,000 worldwide samples. Wellcome Open Res. 6, 42 (2021).

49. Zhu, S. J., Almagro-Garcia, J. & McVean, G. Deconvolution of multiple infections in Plasmodium falciparum from high throughput sequencing data. Bioinforma. Oxf. Engl. 34, 9–15 (2018).

50. Zhu, S. J. et al. The origins and relatedness structure of mixed infections vary with local prevalence of P. Falciparum malaria. eLife 8, e40845 (2019).

51. Cui, L. et al. Malaria in the Greater Mekong Subregion: Heterogeneity and complexity. Acta Trop. 121, 227–239 (2012).

52. Kelleher, J. et al. Inferring whole-genome histories in large population datasets. Nat. Genet. 51, 1330–1338 (2019).

53. Haller, B. C. & Messer, P. W. SLiM 3: Forward Genetic Simulations Beyond the Wright–Fisher Model. Mol. Biol. Evol. 36, 632–637 (2019).

54. Harris, D. N. et al. Evolutionary genomic dynamics of Peruvians before, during, and after the Inca Empire. Proc. Natl. Acad. Sci. U. S. A. 115, E6526–E6535 (2018).

55. Wesolowski, A. et al. Mapping malaria by combining parasite genomic and epidemiologic data. BMC Med. 16, 190–190 (2018).

56. Newman, M. E. J. & Girvan, M. Finding and evaluating community structure in networks. Phys. Rev. E Stat. Phys. Plasmas Fluids Relat. Interdiscip. Top. 69, 026113 (2004).

57. Csardi, G. & Nepusz, T. The igraph software package for complex network research. InterJournal Complex Syst. 1695, 1–9 (2006).

58. Rosvall, M., Axelsson, D. & Bergstrom, C. T. The map equation. Eur. Phys. J. Spec. Top. 178, 13–23 (2009).

59. Schaffner, S. F., Taylor, A. R., Wong, W., Wirth, D. F. & Neafsey, D. E. HmmIBD: Software to infer pairwise identity by descent between haploid genotypes. Malar. J. 17, 10–13 (2018).

60. Koenderink, J. B., Kavishe, R. A., Rijpma, S. R. & Russel, F. G. M. The ABCs of multidrug resistance in malaria. Trends Parasitol. 26, 440–446 (2010).

61. Tindall, S. M. et al. Heterologous Expression of a Novel Drug Transporter from the Malaria Parasite Alters Resistance to Quinoline Antimalarials. Sci. Rep. 8, 2464 (2018).

62. Martin, R. E. & Kirk, K. The Malaria Parasite’s Chloroquine Resistance Transporter is a Member of the Drug/Metabolite Transporter Superfamily. Mol. Biol. Evol. 21, 1938–1949 (2004).

63. Kim, J. et al. Structure and drug resistance of the Plasmodium falciparum transporter PfCRT. Nature 576, 315–320 (2019).

64. Brooks, D. R. et al. Sequence variation of the hydroxymethyldihydropterin pyrophosphokinase: Dihydropteroate synthase gene in lines of the human malaria parasite, Plasmodium falciparum, with differing resistance to sulfadoxine. Eur. J. Biochem. 224, 397–405 (1994).

65. Miotto, O. et al. Genetic architecture of artemisinin-resistant Plasmodium falciparum. Nat. Genet. 47, 226–234 (2015).

66. Kümpornsin, K. et al. Origin of Robustness in Generating Drug-Resistant Malaria Parasites. Mol. Biol. Evol. 31, 1649–1660 (2014).

67. Takala-Harrison, S. et al. Genetic loci associated with delayed clearance of plasmodium falciparum following artemisinin. Proc. Natl. Acad. Sci. U. S. A. 110, 240–245 (2013).

68. Takala-Harrison, S. et al. Independent emergence of artemisinin resistance mutations among Plasmodium falciparum in Southeast Asia. J. Infect. Dis. 211, 670–9 (2015).

69. Birnbaum, J. et al. A Kelch13-defined endocytosis pathway mediates artemisinin resistance in malaria parasites. Science 367, 51–59 (2020).

70. Josling, G. A. et al. Dissecting the role of PfAP2-G in malaria gametocytogenesis. Nat. Commun. 11, 1503 (2020).

71. Ye, R., Zhang, Y. & Zhang, D. Evaluations of candidate markers of dihydroartemisinin-piperaquine resistance in Plasmodium falciparum isolates from the China–Myanmar, Thailand–Myanmar, and Thailand–Cambodia borders. Parasit. Vectors 15, 130 (2022).

72. Hubert, L. & Arabie, P. Comparing partitions. J. Classif. 2, 193–218 (1985).

73. Bamshad, M. & Wooding, S. P. Signatures of natural selection in the human genome. Nat. Rev. Genet. 4, 99–110 (2003).

74. Santiago, E. et al. Recent Demographic History Inferred by High-Resolution Analysis of Linkage Disequilibrium. Mol. Biol. Evol. 37, 3642–3653 (2020).

75. Barbato, M., Orozco-terWengel, P., Tapio, M. & Bruford, M. W. SNeP: A tool to estimate trends in recent effective population size trajectories using genome-wide SNP data. Front. Genet. 6, (2015).

76. Nadachowska-Brzyska, K., Konczal, M. & Babik, W. Navigating the temporal continuum of effective population size. Methods Ecol. Evol. 13, 22–41 (2022).

77. Novo, I., Santiago, E. & Caballero, A. The estimates of effective population size based on linkage disequilibrium are virtually unaffected by natural selection. PLOS Genet. 18, e1009764 (2022).

78. Nkhoma, S. C. et al. Co-transmission of Related Malaria Parasite Lineages Shapes Within-Host Parasite Diversity. Cell Host Microbe 27, 93-103.e4 (2020).

79. Miotto, O. et al. Emergence of artemisinin-resistant Plasmodium falciparum with kelch13 C580Y mutations on the island of New Guinea. PLOS Pathog. 16, e1009133 (2020).

80. Johri, P., Charlesworth, B. & Jensen, J. D. Toward an Evolutionarily Appropriate Null Model: Jointly Inferring Demography and Purifying Selection. Genetics 215, 173–192 (2020).

81. Tang, J., Huang, M., He, S., Zeng, J. & Zhu, H. Uncovering the extensive trade-off between adaptive evolution and disease susceptibility. Cell Rep. 40, 111351 (2022).

82. Hartfield, M., Bataillon, T. & Glémin, S. The Evolutionary Interplay between Adaptation and Self-Fertilization. Trends Genet. 33, 420–431 (2017).

83. Haller, B. C., Galloway, J., Kelleher, J., Messer, P. W. & Ralph, P. L. Tree-sequence recording in SLiM opens new horizons for forward-time simulation of whole genomes. Mol. Ecol. Resour. 19, 552–566 (2019).

84. Baumdicker, F. et al. Efficient ancestry and mutation simulation with msprime 1.0. Genetics 220, iyab229 (2022).

85. Tang, K., Naseri, A., Wei, Y., Zhang, S. & Zhi, D. Open-source benchmarking of IBD segment detection methods for biobank-scale cohorts. GigaScience 11, giac111 (2022).

86. MalariaGEN et al. Pf7: an open dataset of Plasmodium falciparum genome variation in 20,000 worldwide samples. Wellcome Open Res. 8, 22 (2023).

87. Auburn, S. et al. Characterization of Within-Host Plasmodium falciparum Diversity Using Next-Generation Sequence Data. PLOS ONE 7, e32891 (2012).

88. Lee, S. & Bahlo, M. Moimix: An R package for assessing clonality in high-throughput sequencing data. Moimix R Package Assess. Clonality High-Throughput Seq. Data (2016).

89. Browning, B. L., Zhou, Y. & Browning, S. R. A One-Penny Imputed Genome from Next-Generation Reference Panels. Am. J. Hum. Genet. 103, 338–348 (2018).

90. Kelleher, J., Thornton, K. R., Ashander, J. & Ralph, P. L. Efficient pedigree recording for fast population genetics simulation. PLoS Comput. Biol. 14, 1–21 (2018).

91. Bopp, S. E. R. et al. Mitotic Evolution of Plasmodium falciparum Shows a Stable Core Genome but Recombination in Antigen Families. PLOS Genet. 9, e1003293 (2013).

92. Susomboon, P. et al. Differences in genetic population structures of Plasmodium falciparum isolates from patients along Thai-Myanmar border with severe or uncomplicated malaria. Malar. J. 7, 212 (2008).

93. Joy, D. A. et al. Early Origin and Recent Expansion of Plasmodium falciparum. Science 300, 318–321 (2003).

94. Carrasquilla, M. et al. Resolving drug selection and migration in an inbred South American Plasmodium falciparum population with identity-by-descent analysis. PLOS Pathog. 18, e1010993 (2022).

95. Mathieu, L. C. et al. Local emergence in Amazonia of Plasmodium falciparum k13 C580Y mutants associated with in vitro artemisinin resistance. eLife 9, e51015 (2020).

96. Wright, S. Coefficients of Inbreeding and Relationship. Am. Nat. 56, 330–338 (1922).

97. Seabold, S. & Perktold, J. Statsmodels: Econometric and Statistical Modeling with Python. Proc. 9th Python Sci. Conf. 92–96 (2010) doi:10.25080/Majora-92bf1922-011.

