## Supplementary Figures and Tables for "Strong Positive Selection Biases Identity-By-Descent-Based Inferences of Recent Demography and Population Structure in *Plasmodium falciparum*"

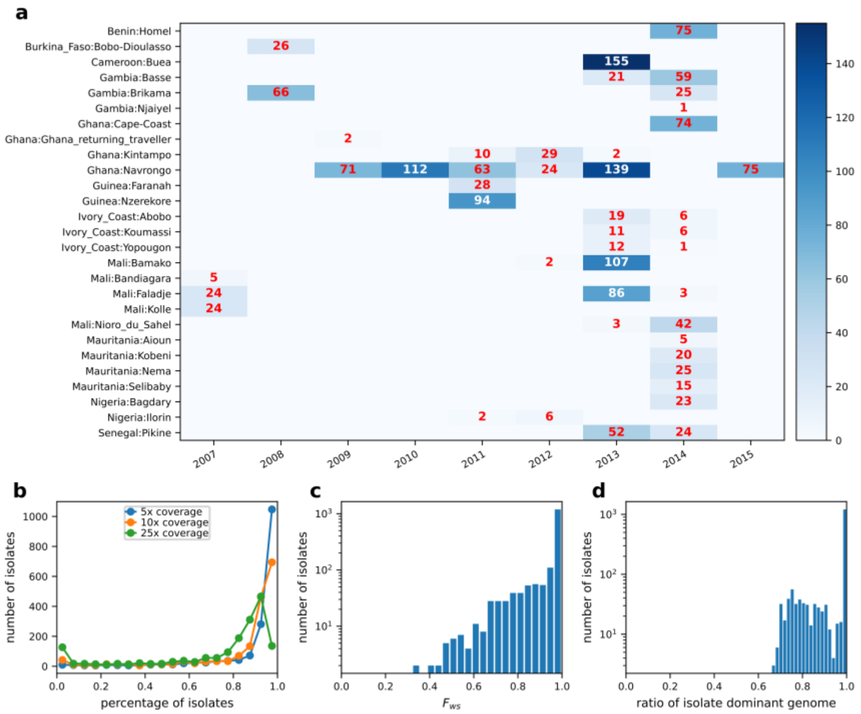


**Figure S1. Summary of samples and WGS data for West Africa. a**, Sampling location and time distribution of 2,611 analyzable samples. **b**, Distribution of genome fractions covered by at least 5, 10, and 25 sequence reads of all collected WGS samples from SEA. **c**, Distribution of the multiplicity of infection metric $F_{\text{ws}}$ in WGS samples passed quality control. **d**, Distribution of ratios of dominant genomes in WGS samples passed quality control (determined by dEploid).

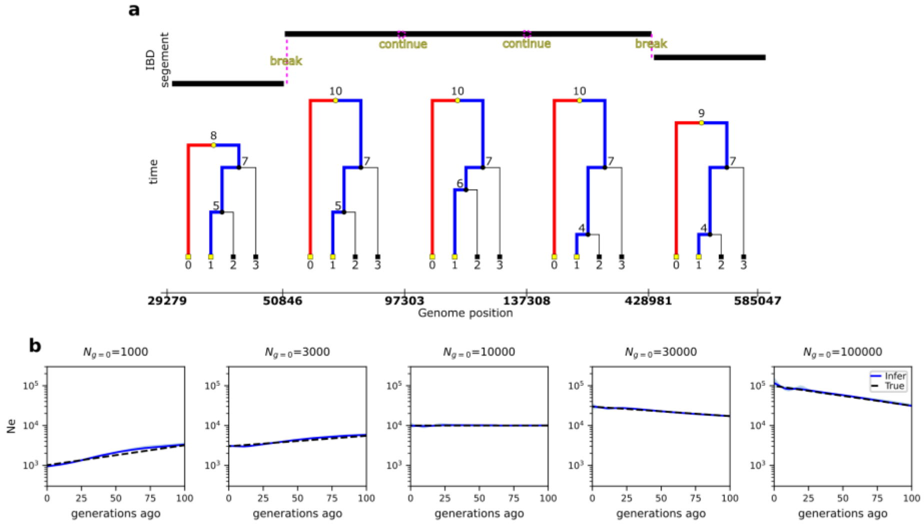


**Figure S2. True IBD inference and its validation in IBD-based $N_e$ estimation. a**, Schematic of true IBD inference process in `tskibd`. For each genealogical tree along the chromosome (low panel), the most recent common ancestor (MRCA) of a pair of genomes (for instance the two with IDs 0 and 1) can be found by tracing backward (red path for genome 0 and blue path for genome 1) until the two paths meet. A theoretical IBD segment is defined by the MRCA (the origin of IBD ) and tree span (the start and end of the IBD segment). Moving along the tree sequence, the IBD segment may break or continue at the tree junction depending on whether recombination happened to this ancestral segment (upper panel). Breaks and continuity can be determined by comparing MRCA node IDs between two nearby sampled trees. To match with most other IBD inference tools, we only include long IBD segments (no shorter than 2 centimorgans) in the output. **b**, True IBD (inferred via `tskibd`)-based $N_e$ estimates for the recent 100 generations are consistent with parameter population size in neutral simulations under different demographic patterns (left two: exponential decrease; middle: constant N; right two: exponential growth).

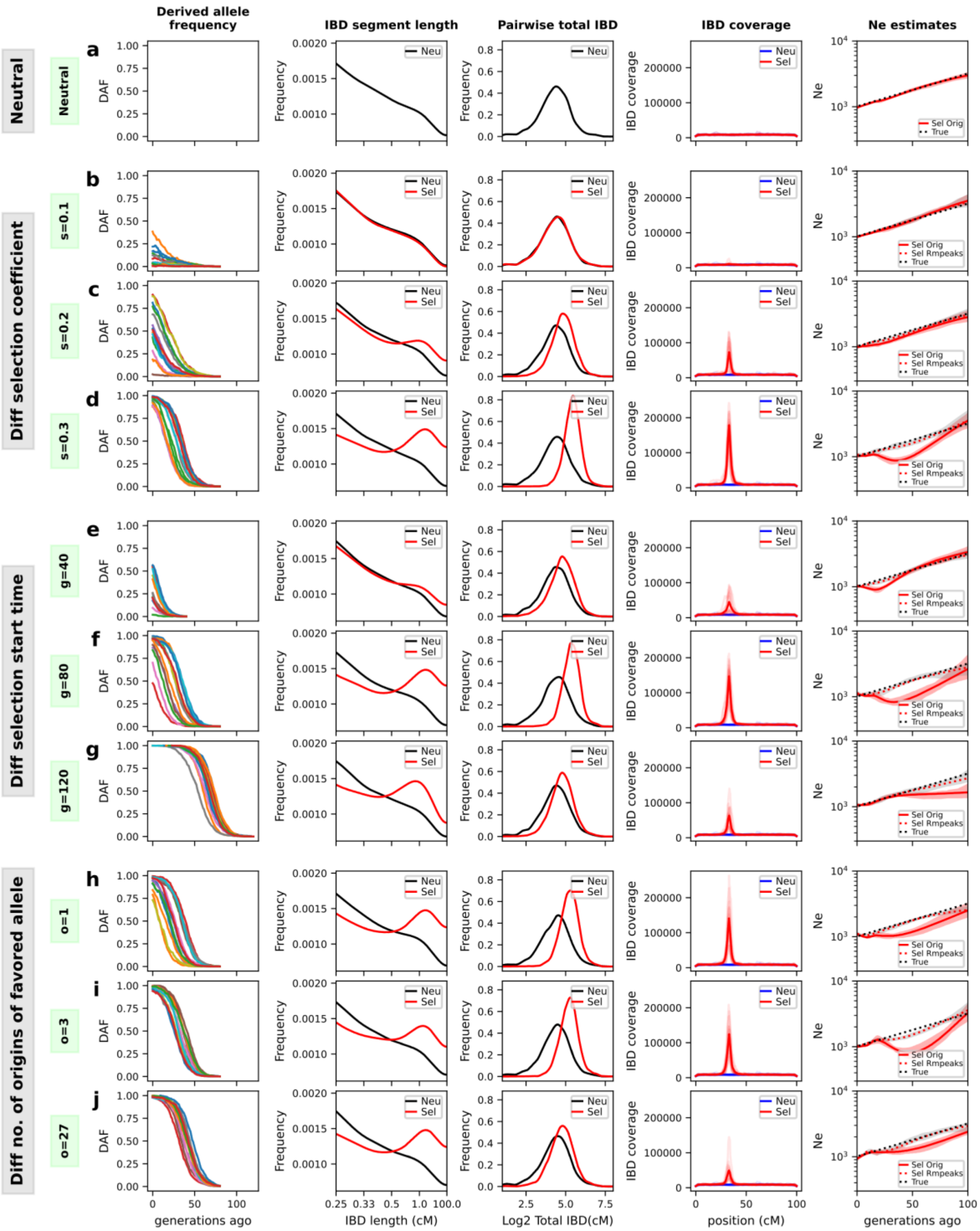


**Figure S3. Extended simulations of selection effect on IBD distribution and $N_e$ estimation.** Different rows are data from different simulation cases, including neutral (**a**), varying selection coefficients (**b-d**), varying selection starting time (**e-g**), and varying numbers of origins of the favored allele (**h-j**). Note that **(a)** x axis (IBD length *l*, bottom) uses a custom scale so that the estimated TMRCA (50/*l*, top) is in a linear scale. Shorter IBD segments (<2 cM) were included to cover the more distant past (>25 generations ago). A length of 0.25, 0.33, 0.5, 1.0 and 100 cM corresponds to a TMRCA of 200, 150, 100, 50 and 0.5 generations ago, respectively. Column 1 are frequency trajectories of the favored alleles on each of the 14 chromosomes. Column 2 shows the IBD length distribution for two types of IBD segments (red: selection simulations, back: neutral simulations). Column 3 shows the distribution of pairwise genome-wide total IBD sharing. Column 4 shows IBD coverage across the chromosome with each color representing 1/14 chromosomes. Column 5 is IBD enrichment profiling (IBD coverage). Column 5 shows$N_e$ estimates and parameter population size. Default parameters: s = 0.3, g = 80, o = 1.

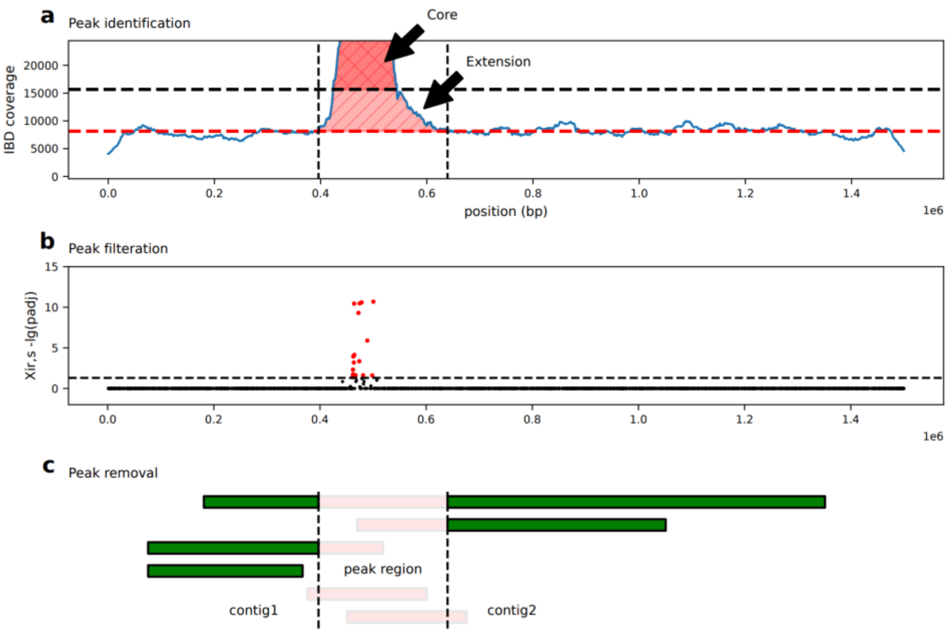


**Figure S4. Schematic for IBD peak region identification and removal. a**, A peak region is a combination of a core region and extension regions. The core region is determined by a threshold of two standard deviations from the chromosomal mean after 5% trimming. This is flanked by extension regions that extend up to the chromosomal median. **b**, $X_{\mathrm{iR},s}$ scan. Red dots (hits) indicate SNPs with adjusted p-value < 0.05 based on IBD-based $X_{\mathrm{iR},s}$ test. Peaks that contain at least one hit are kept and treated as validated peaks. **c**, IBD segments between a pair of isolates (represented by colored bars) are split when they overlap with the peak region (between dotted lines). The segment parts that fall within the peak region (shown in pink) are discarded. This creates a region of IBD emptiness, and the remaining portion of the chromosome is split into contigs containing IBD segments. These contigs are treated as separate chromosomes for `IBDNe` inference.

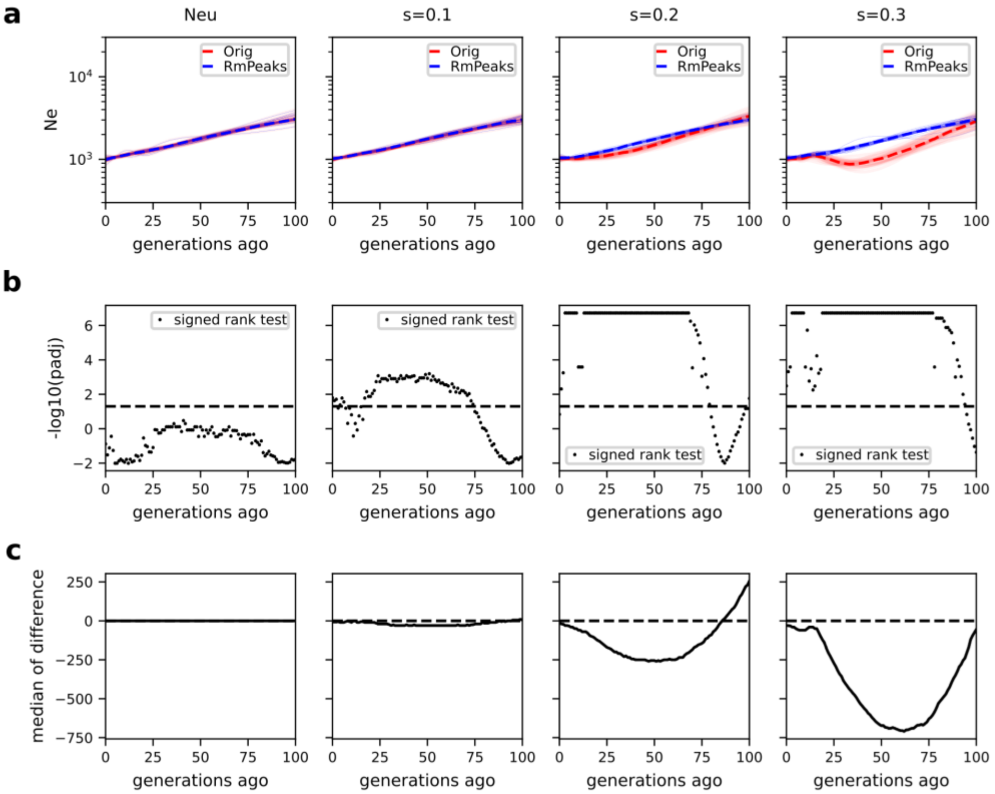


**Figure S5. Validation of positive selection effects on $N_e$ estimates with replicated simulations. a**, $N_e$ trajectories. Solid lines are $N_e$ estimates from 30 replicated simulations Dashed lines are the corresponding medians for each generation. **b**, *p* values of Signed rank test per time point (generation). The horizontal dashed line represents thresholds for significance for the adjusted (Bonferroni method) p values. **c**, Difference of medians per time point between estimates of original and peak-removed IBD segments.

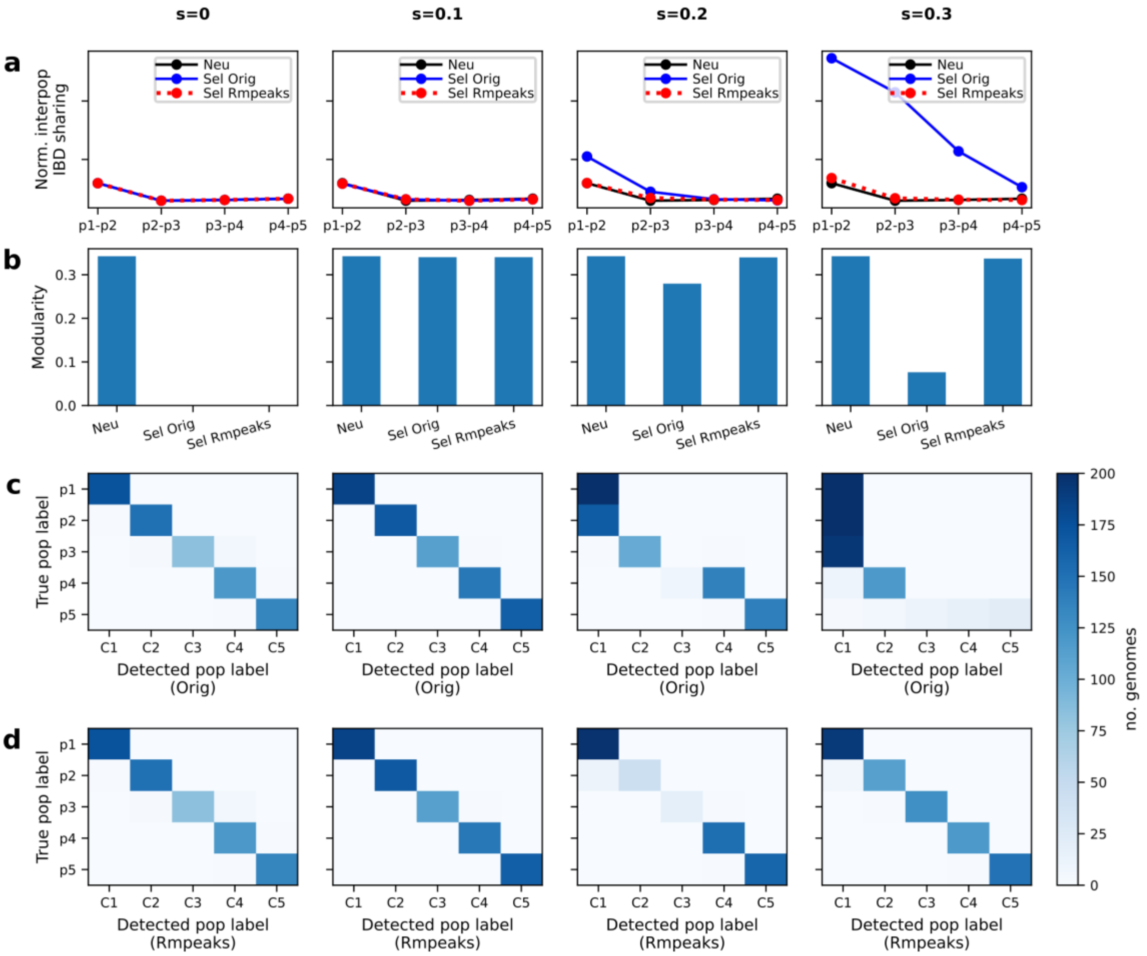


**Figure S6. The effect of positive selection on the IBD-based population structure inference (extending Fig. 3). a**, Schematic of the one-dimensional stepping-stone model. 5 subpopulations were split from an ancestral population. There is symmetrical migration between adjacency subpopulations. A favored allele was introduced in into the deme from one side of the chain and spread to the other side. **b**, Average frequency trajectory of favored alleles (on each chromosome) in different subpopulations (p1 - p5). **c**, Heatmap of pairwise genome-wide total IBD under neutral, selection (s = 0.3), and selection with peaks removed scenarios. Rows and columns are ordered by true population labels. **d**, Normalized inter-population IBD sharing between nearby demes along the stepping stone chain. From left to right are simulations with different coefficients s = 0, s = 0.1, s = 0.2, and s = 0.3. **e**, Modularity of IBD networks with respect to the true population labels under different selection coefficients (neutral, s = 0.1, s = 0.2, and s = 0.3) and IBD processing conditions (before and after removing IBD peaks). **f-g**, IBD network InfoMap community detection before (**f**) and after (**g**) removing IBD peaks. For each subplot, columns are detected cluster (community) labels and rows are true population labels. The color of each block represents the number of isolates with the given true and detect labels.

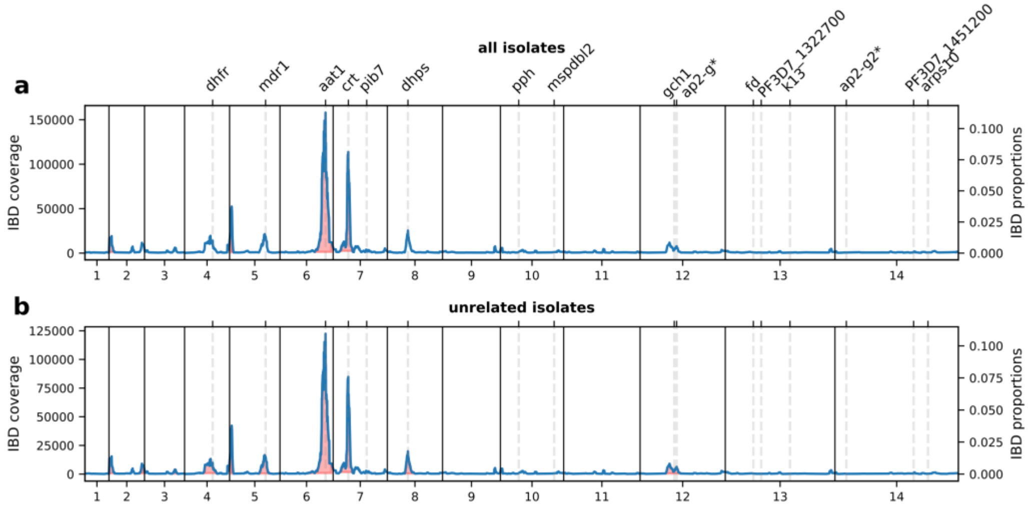


**Figure S7. IBD coverage profile of all and unrelated Pf isolates in WAF (in comparison with SEA, Fig. 1). a**, IBD coverage of all samples in WAF. Labels on the top indicate the center of genes of known or putative drug resistance or that are under selection for sexual commitment (*). **b**, IBD coverage of unrelated samples in WAF. Annotations in A are shared with B.

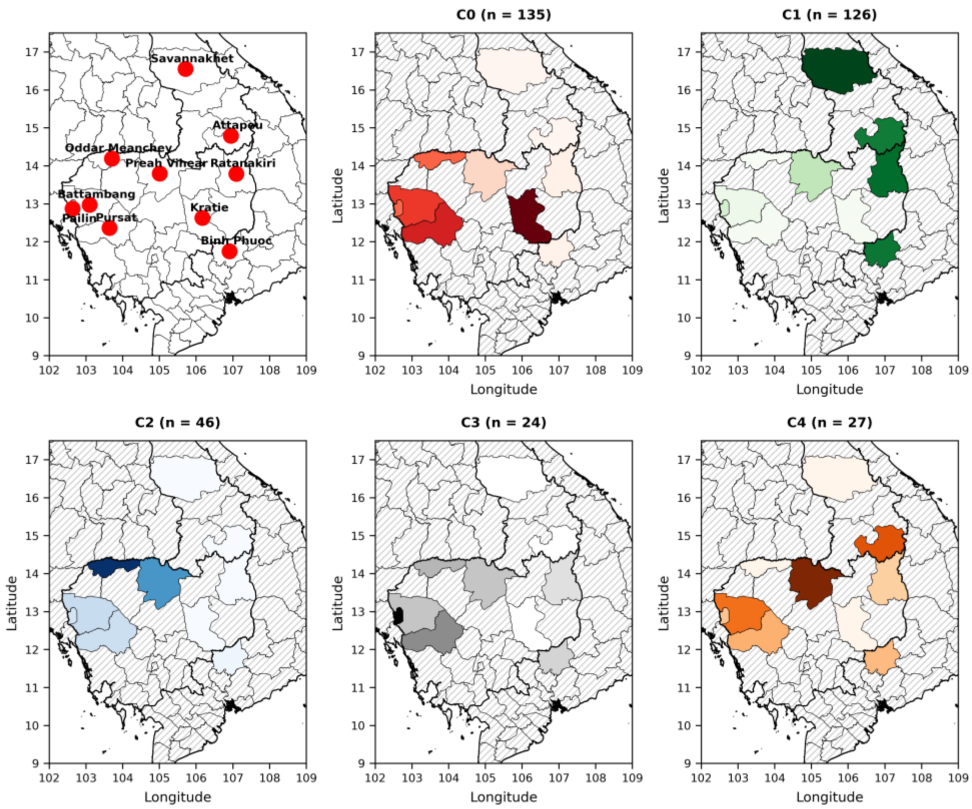


**Figure S8. The enrichment of the largest 5 SEA *Pf* communities in provinces.** The top left panel shows the province labels. The rest of the panels show the enrichment of a given SEA community in different provinces. The enrichment is normalized within each province (the total enrichment of these 5 communities is 100% per province). The darkness of color represents the extent of enrichment of the community in a province. Only provinces with at least 10 samples assigned to these 5 communities are shown. Provinces with hatches do not have enough data.

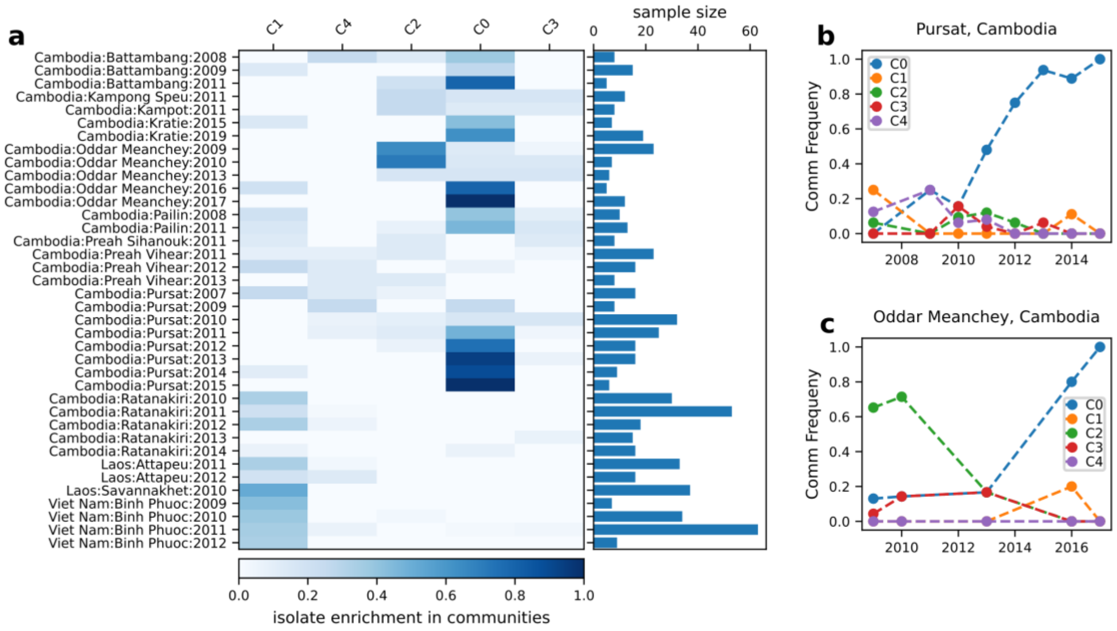


**Figure S9. The enrichment of the largest 5 SEA communities in each region/year combination. a**, Heatmap showing enrichment of top communities in each region-year. The color in the heatmap represents the percentage of samples in a region-year combination (rows) that are assigned to a community (columns). The bar plot on the right of the heatmap shows the number of unrelated samples that are from a region-year (rows). **b-c**. Dynamics in community enrichments in Pursat and Oddar Meanchey provinces in Cambodia.

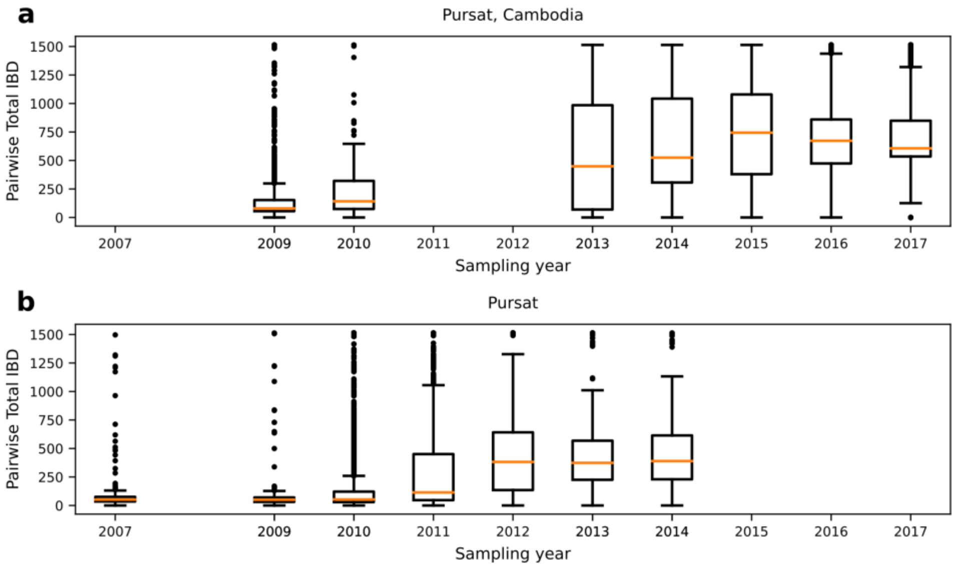


**Figure S10. Dynamics of pairwise genome-wide total IBD sharing over years. a**, Boxplot showing pairwise total IBD in the Oddar Meanchey province of Cambodia. **b**, Boxplot showing pairwise total IBD in the Pursat province of Cambodia. Each time point represents a year during which at least 30 isolates were collected in the given province (including both related and unrelated isolates). The lower and upper edges of the box are the first and third quartile of the data per time point. The center line indicates the median values. The whiskers extend from the box by 1.5x the inter-quartile range. Flier points are those beyond the whisker range.

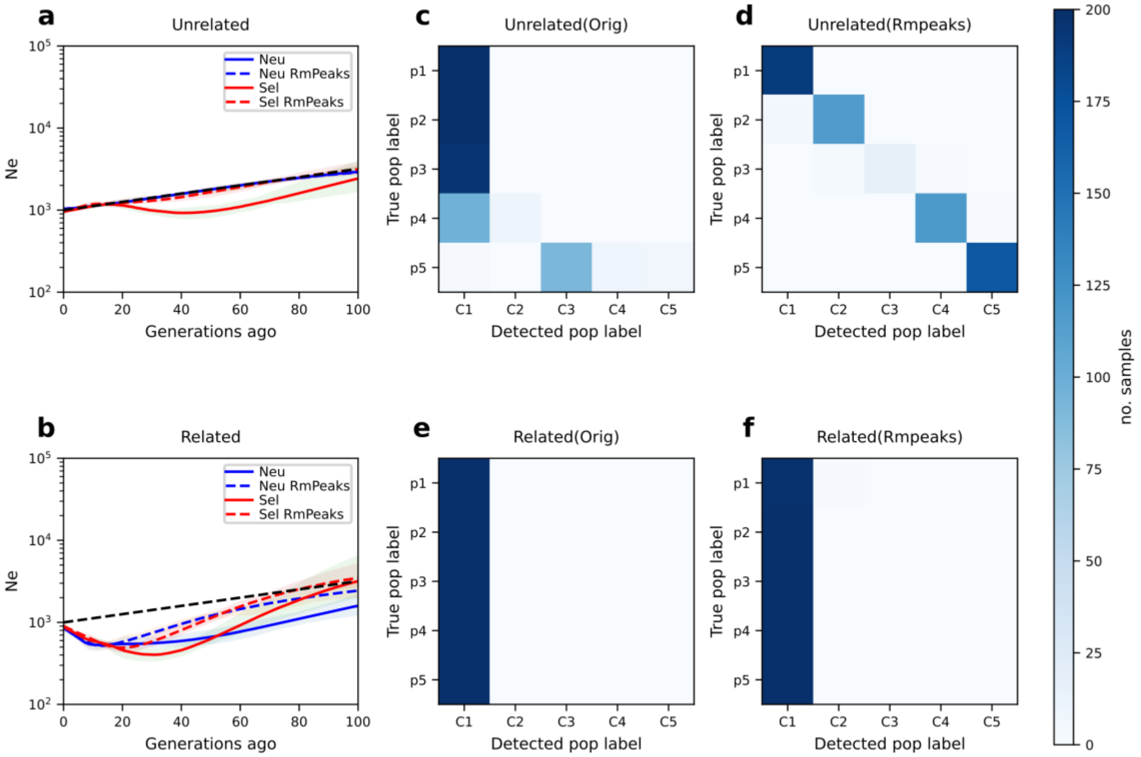


**Figure S11. Positive selection effects can be hidden due to high relatedness in simulation. a-b**, IBD-based $N_e$ estimates for simulated data without (**a**) or with (**b**) incorporating high relatedness. Solid and dashed lines are estimates based on IBD segments before and after removing peaks, respectively. **c-f**, Heatmaps of confusion matrix of IBD network community detection in simulated data without (**c-d**) or with (**e-f**) the incorporating high relatedness. Middle panels: before removing IBD peaks; right panels: after removing IBD peaks. For each heatmap, the top 5 communities (columns) are included and plotted against the true classes (rows).

**Table S1. The effects of positive selection on IBD-based population assignments and the benefits of selection correction are selection-strength dependent (Simulation).** For each parameter (neutral, s = 0.1, s=0.2, and s = 0.3), 30 independent simulations were performed. Adjusted Rand Scores were presented as mean ± standard deviation.

| | AdjRand (Orig) | AdjRand (Rmpeaks) | PvaluePairedTtest |
|---|---|---|---|
| Group | | | |
| Neu | 0.603 ± 0.061 | 0.603 ± 0.061 | 1.0 |
| s=0.1 | 0.622 ± 0.066 | 0.620 ± 0.062 | 0.336 |
| s=0.2 | 0.488 ± 0.072 | 0.629 ± 0.079 | 0.000 |
| s=0.3 | 0.124 ± 0.035 | 0.559 ± 0.084 | 0.000 |

**Table S2. Estimating uncertainty in population structure inference for empirical data based on Jackknife resampling over chromosomes**. The top table shows details of community detection results in each Jackknife sampling. The bottom table shows whether selection correction significantly changes IBD-based community detection in two transmission settings. Orig/Rmpeaks: results for IBD segments before and after removing peaks. NCommu: total number of communities inferred. TopComm: the size (number of samples) in the largest 5 communities. AdjRand: adjust rand score based on samples that exist in the top 5 communities of both Orig and Rmpeaks results. RmChr: the chromosome number was removed in Jackknife resampling. Q3, the third quartile; IQR: interquartile range, Q3 - Q1.

| | | **Orig** | | **Rmpeaks** | | | |
|---|---|---|---|---|---|---|---|
| Region | N | NComm | TopComm | NComm | TopComm | AdjRnd | RmChr |
| SEA | 701 | 72 | 134,127,44,28,23 | 81 | 141,86,38,26,25 | 0.933 | 12 |
| SEA | 701 | 73 | 137,129,46,24,24 | 84 | 137,83,31,25,19 | 0.960 | 6 |
| SEA | 701 | 73 | 141,139,44,26,23 | 83 | 138,75,41,23,21 | 0.972 | 5 |
| SEA | 701 | 74 | 139,126,45,27,24 | 83 | 139,72,41,24,20 | 0.980 | 3 |
| SEA | 701 | 74 | 139,128,43,24,23 | 80 | 135,80,31,23,21 | 0.973 | 7 |
| SEA | 701 | 74 | 139,135,44,27,23 | 83 | 141,90,28,24,20 | 0.952 | 8 |
| SEA | 701 | 74 | 140,132,42,25,24 | 81 | 137,92,43,25,24 | 0.982 | 4 |
| SEA | 701 | 75 | 134,115,45,24,24 | 87 | 131,92,31,25,21 | 0.979 | 11 |
| SEA | 701 | 75 | 134,119,45,27,26 | 83 | 131,85,30,28,24 | 0.958 | 13 |
| SEA | 701 | 75 | 135,126,46,27,24 | 85 | 136,79,43,25,23 | 0.956 | None |
| SEA | 701 | 75 | 139,131,43,25,25 | 86 | 139,66,41,23,21 | 0.946 | 2 |
| SEA | 701 | 76 | 134,132,44,24,24 | 86 | 138,69,31,26,23 | 0.971 | 1 |
| SEA | 701 | 76 | 143,117,40,24,24 | 85 | 137,57,32,27,22 | 0.985 | 14 |
| SEA | 701 | 77 | 136,136,42,24,23 | 84 | 133,96,40,25,20 | 0.982 | 10 |
| SEA | 701 | 78 | 137,127,43,24,23 | 85 | 134,80,42,25,24 | 0.974 | 9 |
| WAF | 1496 | 67 | 1238,25,16,16,12 | 189 | 57,41,37,36,34 | 0.092 | 12 |
| WAF | 1496 | 69 | 1237,25,17,15,14 | 184 | 53,40,39,38,34 | 0.097 | 13 |
| WAF | 1496 | 69 | 1238,24,16,14,12 | 102 | 1035,55,29,26,23 | 0.523 | 10 |
| WAF | 1496 | 71 | 1228,24,17,16,14 | 168 | 213,56,36,34,33 | 0.140 | 3 |
| WAF | 1496 | 71 | 1229,25,17,16,11 | 175 | 59,39,36,35,33 | 0.095 | 8 |
| WAF | 1496 | 71 | 1233,25,19,14,12 | 185 | 58,40,39,38,34 | 0.091 | 14 |
| WAF | 1496 | 73 | 1223,25,18,18,11 | 174 | 62,47,41,39,36 | 0.081 | 5 |
| WAF | 1496 | 74 | 1221,25,17,16,12 | 183 | 54,42,38,37,33 | 0.087 | 1 |
| WAF | 1496 | 74 | 1225,24,17,16,12 | 180 | 57,46,39,38,35 | 0.074 | 2 |
| WAF | 1496 | 74 | 1230,24,16,13,12 | 182 | 63,40,36,36,33 | 0.136 | 9 |
| WAF | 1496 | 74 | 1230,24,17,16,12 | 186 | 55,46,44,38,35 | 0.074 | 11 |
| WAF | 1496 | 75 | 1219,25,18,16,12 | 190 | 60,35,35,33,31 | 0.169 | 7 |
| WAF | 1496 | 75 | 1222,25,17,16,11 | 175 | 61,49,42,37,35 | 0.085 | None |
| WAF | 1496 | 76 | 1223,25,17,16,12 | 176 | 75,60,39,37,36 | 0.089 | 4 |
| WAF | 1496 | 83 | 1170,27,25,19,11 | 97 | 1093,31,27,22,12 | 0.913 | 6 |

| Region | Q3 + 1.5 IQR | Significant |
|---|---|---|
| SEA | 1.01 | False |
| WAF | 0.22 | True |
